# Loss of *AUXIN RESPONSE FACTOR 4* function alters plant growth, stomatal function and improves tomato tolerance to salinity and osmotic stress

**DOI:** 10.1101/756387

**Authors:** Sarah Bouzroud, Karla Gasparini, Guojian Hu, Maria Antonia Machado Barbosa, Bruno Luan Rosa, Mouna Fahr, Najib Bendaou, Mondher Bouzayen, Agustin Zsögön, Abdelaziz Smouni, Mohamed Zouine

## Abstract

Auxin controls multiple aspects of plant growth and development. However, its role in stress responses remains poorly understood. Auxin acts on the transcriptional regulation of target genes, mainly through Auxin Response Factors (*ARF*). This study focuses on the involvement of *SlARF4* in tomato tolerance to salinity and osmotic stress. Using a reverse genetic approach, we found that the antisense down-regulation of *SlARF4* promotes root development and density, increases soluble sugars content and maintains chlorophyll content at high levels under stress conditions. Furthermore, *ARF4*-as displayed higher tolerance to salt and osmotic stress through reduced stomatal conductance coupled with increased leaf relative water content and ABA content under normal and stressful conditions. This increase in ABA content was correlated with the activation of ABA biosynthesis genes and the repression of ABA catabolism genes. *cat1, Cu/ZnSOD* and *mdhar* genes were up-regulated in *ARF4*-as plants which can result in a better tolerance to salt and osmotic stress. A CRISPR/Cas9 induced *SlARF4* mutant showed similar growth and stomatal responses as *ARF4*-as plants, which suggest that *arf4-cr* can tolerate salt and osmotic stresses. Our data support the involvement of *ARF4* as a key factor in tomato tolerance to salt and osmotic stresses and confirm the use of CRISPR technology as an efficient tool for functional reverse genetics studies.

## 1. Introduction

Auxin is a key regulator of many plant growth and development processes throughout the plant life cycle [1]. Auxins regulate diverse cellular and developmental responses in plants via three types of transcriptional regulators, auxin response factors (ARFs), Aux/IAA and TOPLESS (TPS) proteins [2]. ARFs play a key role in regulating the expression of auxin response genes. 23 *ARF* genes have been isolated from *Arabidopsis thaliana* [3]. Most of them share a structure with conserved domains: an N-terminal DNA-binding domain (DBD), a variable central transcriptional regulatory region (MR), which can function as an activator or repressor domain, and a carboxy-terminal dimerization domain (CTD) that contributes to the formation of either ARF/ARF homo- and hetero-dimers or ARF/Aux/IAA hetero-dimers [4]. Since *ARF1*, the first *Arabidopsis ARF* gene, was cloned and its function investigated [5], the *ARF* gene family has been identified and well characterized in many crop species, including tomato (*Solanum lycopersicum*), maize (*Zea mays*), rice (*Oryza sativa*), poplar (*Populus trichocarpa*), Chinese cabbage (*Brassica rapa*), banana (*Musa* sp.) and physic nut (*Jatropha curcas* L.) [4,6–9].

In tomato, the *ARF* gene family is also involved in the control of many physiological processes. Downregulation of *ARF3* decreased the density of epidermal cells and trichomes in tomato [10]. The *Slarf2A*/*B* mutation leads to a severe fruit ripening inhibition with dramatically reduced ethylene production, while the over-expression of *ARF2A* resulted in a blotchy ripening pattern in fruit as a result of a significant accumulation of ripening-related genes and metabolites [11]. Down-regulation of *SlARF6* and *SlARF8* in transgenic plants through the overexpression of *miR167a* may lead to floral development defects and female sterility [12]. Transcriptional down-regulation of *ARF4* expression leads to severe leaf curling along the longitudinal axis [13]. The down-regulation of *SlARF4* also resulted in ripening-associated phenotypes such as enhanced firmness, sugar and chlorophyll content leading to dark green fruit and blotchy ripening [13,14]. This phenotype was also reported in *SlARF10* gain of function mutant [15].

*ARF* genes also seem to be implicated in plant responses to environmental stresses. It has been reported that both *OsARF11* and *OsARF15* show differential expression in salt stress conditions, suggesting that they are involved in the rice response to stress [6]. In banana, the expression of many *ARF* genes was altered in response to salinity and osmotic stresses [8]. We have previously shown that, in tomato, expression levels of many *SlARF* genes are responsive to a wide range of abiotic stress conditions [16]. Interestingly, the *SlARF4* regulatory region was enriched in cis-*acting* elements specific to salinity and water deficit response [16]. Here, we extended this observation to plants subjected to osmotic, salinity and osmotic stress. We investigated whether loss of *SlARF4* function in tomato could have an impact on plant growth and function under water deficit, high salinity or osmotic stress conditions. By assessing morphological, anatomical, physiological and molecular analyses, we provide evidence for the involvement of *SlARF4* in tomato response to osmotic, salinity and osmotic stress. We discuss the potential role of elements of the auxin signaling network in responses to environmental stresses.

## 2. Material and Methods

### 2.1. Plant material

Wild-type tomato (*Solanum lycopersicum*, L.) cv Micro-Tom (WT), and the following *AUXIN RESPONSE FACTOR 4* lines were used in this study: an *ARF4* RNAi antisense silenced line (*ARF4*-as), and a reporter line with the β-glucoronidase (GUS) gene driven by the native *ARF4* promoter region (pARF4∷GUS). *ARF4*-as plant lines were previously generated and well-characterized by Sagar et al. (2014) [14]. The most representative plant line was chosen.

### 2.2. Generation of SlARF4-Crispr (arf4-cr) plants

Tomato *SlARF4*-crispr (*arf4-cr*) plants were obtained by *Agrobacterium tumefasciens* mediated genetic transformation according to Wang et al. (2005) [17]. CRISPR/Cas9 mutant lines were generated following previous standard protocol [18]. Two single-guide (sg) RNAs in the Solyc11g069190 coding sequence were designed using the CRISPR-P server (http://cbi.hzau.edu.cn/cgi-bin/CRISPR) [19]. The sgRNA sequences are AATGGAGGTCACACCAGAG and GGAACTGAAAAGCCACCAT, aiming to create a 49bp deletion in the DBD domain of *SlARF4* gene. The sgRNAs were first cloned in the Level 1 vectors pICH47751 and pICH47761 driven by the *Arabidopsis* U6 promoter, respectively. The Level 1 constructs pICH47732-NOSpro∷NPTII, pICH47742-35S:Cas9, pICH47751-AtU6pro:sgRNA1, and pICH47761-AtU6∷sgRNA2 were then assembled into the Level 2 destination vector pAGM4723. For genotyping of the transgenic lines, genomic DNA was extracted using ReliaPrep™ gDNA Tissue Miniprep System (Promega). The CRISPR/Cas9-positive lines were identified by PCR, then further genotyped for mutations using the primers (Fwd: ACATGGTTTCACTGTAAAGGGATCT and Rev: CTGGCCTGAAAGAAAAGCATCAAA) spaning the two sgRNAs target sequences by PCR and Sanger sequencing of ARF4-PCR products.

### 2.3. Characterization of ARF4-as and arf4-cr plants

#### 2.3.1. Plant growth conditions

Seeds of WT, *ARF4*-as and *arf4-cr* transgenic plants were sown in germination trays with commercial substrate (Tropstrato HT Hortaliças, Vida Verde) and supplemented with 1g L^-1^ 10:10:10 NPK and 4 g L^-1^ dolomite limestone (MgCO_3_ + CaCO_3_). After the emergence of the first pair of true leaves, the seedlings were transferred to plastic pots (350mL) with the same substrate described above, supplemented with 8 g L^-1^ 10:10:10 nitrogen: phosphorus: potassium (N:P:K) and 4 g L^-1^ lime. The experiment was conducted in greenhouse localized at Universidade Federal de Viçosa (642 m asl, 20°45’ S; 42°51’ W), Minas Gerais, Brazil, under semi-controlled conditions (mean temperature of 28°C and 450-500 μmol m^-2^ s^-1^ PAR irradiance). Irrigation was performed twice a day, where each plastic pot received the same volume of water.

#### 2.3.2. Growth analyses

Plant height and internode length were measured 35 days after germination (DAG). At 45 DAG the plants were harvest and divided into leaves, stem and root. Leaf area was measured using leaf area meter (Li-Cor Model 3100 Area Meter, Lincoln, NE, USA). Leaf, stem and root were then packed separately in paper bags and oven dried at 70°C for 72h until they reached constant weight. Shoot and root biomass were measured from the dry weight of leaves, stem and root. Specific Leaf Area (SLA) was determined as described by Hunt (1982) [20].

#### 2.3.3. Microscopy

For anatomical analysis, leaf discs were collected from the medium point of the fifth lateral leaflet and fixed in FAA50 (Formaldehyde, acetic acid and ethanol 50%) for 48 h, and then stored in ethanol 70%. Samples were infiltrated with historesin (Leica Microsystems, Wetzlar, Germany) and cut in cross sections with ∼5 μm (RM2155, Leica Microsystems, Wetzlar, Germany). Leaf cross sections were mounted in water on glass slides and stained with toluidine blue. Histological sections were observed using an optic microscope (AX-70 TRF, Olympus Optical, Tokyo, Japan) and then photographed using digital photo camera (Zeiss AxioCam HRc, Göttinger, Germany). Anatomical features, as leaf thickness and thickness cell layers, were analyzed using Image-Pro Plus® software (version 4.5, Media Cybernetics, Silver Spring, USA).

#### 2.3.4. Measurements of Photosynthetic Parameters

Photosynthetic parameters were performed using an open-flow infrared gas exchange analyzer system (Li 6400XT, Li-Cor, Lincoln, USA) equipped with an integrated fluorescence chamber (Li-6400-40; Li-Cor Inc.). All measurements were made on terminal leaflets of intact and completely expanded leaves. The analysis was conducted under common conditions for photon flux density (1000 μmol m-2 s -1), leaf temperature (25 ± 0.5°C), leaf-to-air vapor pressure difference (16.0 ± 3.0 mbar), air flow rate into the chamber (500 μmol s-1), reference CO_2_ concentration of 400 ppm using an area of 2 cm^2^ in the leaf chamber.

*A*/Ci curves were determined initiated at an ambient CO_2_ concentration of 400 μmol mol^−1^ under a saturating photosynthetic Photon Flux Density (PPFD) of 1000 μmol m^−2^ s ^−1^ at 25°C under ambient O_2_ supply. CO_2_ concentration was decreased to 50 μmol mol^−1^ of air in step changes. Upon the completion of the measurements at low C_a_, C_a_ was returned to 400 μmol mol^−1^ of air to restore the original *A*. Next, CO_2_ was increased stepwise to 1600 μmol mol^−1^ of air. The maximum rate of carboxylation (*V*_*cmax*_), maximum rate of carboxylation limited by electron transport (J_*max*_) and triose-phosphate utilization (TPU) were estimated by fitting the mechanistic model of CO_2_ assimilation proposed by Farquhar et al. (1980)[21]. Corrections for the leakage of CO_2_ into and out of the leaf chamber of the LI-6400 were applied to all gas-exchange data as described by Rodeghiero et al. (2007) [22].

#### 2.3.5. Kinetics of stomatal conductance

The evaluation of stomatal behavior in response to light and CO_2_ variation was performed in WT and *ARF4*-as plants 40 DAG, under laboratory conditions (temperature 22-25°C; relative humidity 50-60% and radiation of 150-200 µmol m^-2^ s^-1^), using an open-flow infrared gas exchange analyzer system (Li 6400XT, Li-Cor, Lincoln, USA). The variation of *g*_s_ was performed on the fifth or sixth fully expanded leaf. For step-change evaluation of *g*_s_ in response to light intensity, leaves were stabilized for 30 min in the dark and then radiation was increased to 1000 µmol m^-2^ s^-1^, and allowed to stabilize for 240 min. Subsequently, light was turned off and observations recorded for another 100 min. The alternation in the amount of light was based on the results presented by. The response of *g*_*s*_ to CO_2_ was performed in a similar manner to those described above for light response. The analysis was performed in a range of 240 minutes, with changes in CO_2_ concentration (400-800-400 µmol CO_2_ m^-2^ s^-1^).

#### 2.3.6. Productivity traits

The productivity parameters were measured in eight plants per genotype (WT, *ARF4*-as and *SlARF4-*crispr). The average fruit weight was determined after individual weighing of each fruit, using a semi analytical balance with a sensitivity of 0.01 g (Shimadzu^®^ AUY220 model). Equatorial and polar diameter was measured using a digital pachymeter (Jomarca *STAINLESS HARDENED*). The determination of the soluble solids content (°Brix) was performed in 10 fruits per plant using a digital temperature-compensated refractometer, model RTD 45 (Instrutherm^®^, São Paulo, SP).

### 2.4. Salt and osmotic stress analyses

#### 2.4.1. Plant growth and stress conditions

WT tomato seeds were sterilized for 10 min in 50% sodium hypochlorite, rinsed four times with sterile distilled water and sown in pots containing peat. Then they were incubated in a culture room with 16h light/ 8h dark photoperiod and 25± 2°C temperature. After 3 weeks, plants were subjected to salt and drought stresses. Salt stress was performed by watering daily the plants with 250mM of NaCl solution. Control plants were daily watered with distilled water. Leaves and roots samples were harvested after 2 and 24 hours of salt stress application. Drought stress was performed on three-week-old plants by water holding for 48 hours and for 5 days. Watering continued normally throughout for control plants. Leave and root samples were collected after drought stress application. Three biological replicates were done for each condition.

#### 2.4.2. RNA extraction

Total RNA was extracted from leaves and roots samples by using The Plant RNeasy extraction kit (RNeasy Plant Mini Kit, Qiagen, Valencia, CA, USA). To remove any residual genomic DNA, the RNA was treated with an RNase-Free DNase according to the manufacturer’s instruction (Ambion® DNA-*free*™DNase). The concentration of RNA was accurately quantified by spectrophotometric measurement and 1µg of total RNA was separated on 2% agarose gel to monitor its integrity. DNase-treated RNA (2μg) was then reverse-transcribed in a total volume of 20μl using the Omniscript Reverse Transcription Kit (Qiagen).

#### 2.4.3. Real time PCR

The real-time quantification of cDNA corresponding to 2µg of total RNA was performed in the ABI PRISM 7900HT sequence detection system using the QuantiTech SYBR Green RT-PCR kit (Qiagen). The Gene-specific primers used are listed in Table S1. The reaction mixture (10µl) contained 2µg of total RNA, 1,2 µM of each primer and appropriate amounts of enzymes and fluorescent dyes as recommended by the manufacturer. Actin gene was used as reference. Real-Time PCR conditions were as follow: 50°C for 2 min, 95°C for 10 min, then 40 cycles of 95°C for 15 s and 60°C for 1 min, and finally one cycle at 95°C for 15 s and 60°C for 15 s. Three independent biological replicates were used for real-time PCR analysis. For each data point, the C_T_ value was the average of C_T_ values obtained from the three biological replicates.

#### 2.4.4. Histochemical analysis of GUS expression

Transgenic plants expressing pARF4∷GUS were generated by *A. tumefaciens*-mediated transformation according to Wang et al. (2005) [17]. For that, PCR was performed on the genomic DNA of tomato Micro-Tom (10 ng.ml^−1^) using specific primers. The corresponding amplified fragment was cloned into the pMDC162 vector containing the GUS reporter gene using Gateway technology (Invitrogen). The cloned *SlARF* promoter was sequenced from both sides using vector primers in order to see whether the end of the promoter is matching with the beginning of the reporter gene. Sequencing results analyses were carried out using the Vector NTI (Invitrogen) and ContigExpress software by referring to *ARF* promoter sequences.

After being surface sterilized, pARF4∷GUS seeds were cultivated in Petri dishes containing half strength Murashige & Skoog medium for 7 days in a growth chamber at 25°C with 16h light/ 8h dark cycle. One week plants were then grown hydroponically for two weeks in Broughton & Dillworth (BD) liquid medium [23]. Three-week-old plants were subjected to salt and osmotic treatment. Salt stress was performed by adding 250 mM of NaCl to the culture medium. After 24 hours of the salt stress application, plants were incubated overnight at 37°C in GUS buffer (3 mM 5-bromo-4-chloro-3-indolyl-β-D-glucuronide [Duchefa Biochemie, Haarlem, The Netherlands], 0.1% v/v Triton X-100 [Sigma, Steinhaim, Germany] 8 mM β-mercaptoethanol, 50 mM Na_2_HPO_4_/NaH_2_PO_4_ [pH 7.2]) then followed by a destaining in 70% EtOH.. Osmotic stress was conducted by adding 15% PEG 20000 to the liquid culture solution. Plants were collected after five days of stress application and were GUS-stained as described above. For each stress condition, control plants were cultivated in BD liquid medium for the same period.

#### 2.4.5. Stress tolerance assays in the transgenic tomato plants

WT and *ARF4*-as seeds were first surface-sterilized for 10 min in 50% sodium hypochlorite, rinsed four times with sterile distilled water. They were then cultured in petri dishes containing half strength Murashige & Skoog (MS) medium for 7 days in a growth chamber at 25° with 16h light/ 8h dark cycle. One week plants were then grown hydroponically in pots containing 1L of BD medium for two weeks in the same growth conditions (25° with 16h light/ 8h dark cycle) [23]. Three weeks plants were then subjected to salt and osmotic stresses. Salt stress was performed by adding 100 mM or 150 mM NaCl to the liquid BD medium. Leaves and roots samples were harvested after 2 weeks of treatment. osmotic treatment was conducted as follows: three weeks plants were subjected to osmotic stress by adding 5% or 15% PEG 20 000 corresponding to final osmotic potentials of −0,09 MPa and - 0,28 MPa to the hydroponic solution for 2 weeks. Meanwhile, the control plants were grown normally in BD medium. Leaves and roots samples were collected after 2 weeks of culture for both stressed and unstressed plants. For each treatment, the hydroponic solution was renewed each 3 days.

##### Determination of shoot and root fresh weights

Shoot and root fresh weights were determined as followed: stressed and unstressed plants from each line were harvested and rinsed thoroughly with distilled water (DW). The plants were blot dried on blotting sheet and cut into shoots and roots. The fresh weight of each part of the plant was measured and the mean of shoots and roots weights was determined based on at least twelve plants per line in each condition.

##### Determination of primary root length and lateral root density

Primary root length was determined was determined as followed. Pictures for each plant were analyzed using ImageJ software in order to determinate the root length. Twelve independent plants from each line under control and stress conditions were used to calculate the mean.

The lateral root number was determined by counting the number of emerging roots and the mean was calculated based on at least twelve plants per line in each condition. The lateral root density (LR density) was determined using the following equation: number of LRs / the length of the root.

##### Determination of chlorophyll content

Determination of chlorophyll content of each line under stressful and normal conditions was performed as described in Bassa et al. (2012) [24]. A 100 mg aliquot of leaves collected from stressed and unstressed plants was weighted and ground with 1 ml of 80% acetone. The liquid obtained was centrifuged for 1 min at 10000 rpm to remove any remaining solid tissue. Samples were analyzed by spectrophotometry at two wavelengths, 645 and 663 nm, using 80% acetone as the blank. The total chlorophyll content was determined using the following equations: Total Chlorophyll Content = 20.2 × Chl a + 8.02 × Chl b (Chl a = 0.999A663 – 0.0989A645 and Chl b = –0.328A663+1.77A645)

##### Determination of soluble sugar content

Total soluble sugar content was determined as in Dubois et al. (1956) [25]. 100 mg of fresh leaves and roots of each sample were homogenized in 5 ml of 80 % ethanol in test tubes. Test tubes were kept in water bath of 80 °C for 1 hour, and sample extracts were transferred to another test tube, and 0.5 ml distilled water and 1 ml of 5 % phenol then added and allowed to incubate for 1 hour. Finally, after 1 hour, 2.5 ml sulphuric acid was added to the test tubes and shaken well on an orbital shaker. Absorbance was read at 485 nm on Spectrophotometer. Sucrose was used as standard.

##### Determination of leaf stomatal conductance

Leaf abaxial stomatal conductance of the youngest fully expanded leaf (usually the fifth leaf, counting from the base) of stressed and well-drained plants of each studied line were determined after two weeks of stress treatment using a hand-held leaf diffusion poromoter (SC-1 LEAF POROMETER, Decagon, USA). For each line, three biological replicates were conducted at each condition.

##### Determination of ABA content

Three weeks ARF4-as and WT plants were exposed to salt and osmotic stress. Leaf samples were taken after 24 hours for salt stress and after 48 hours for osmotic. ABA measurement assays were performed as described Forcat et al. (2008) [26]. Briefly, 110 mg of frozen tissue were extracted at 4°C for 30 min with 400 μl of H_2_O with 10% methanol + 1% acetic acid. The internal standard was 2H6 ABA. The extract was centrifuged at 13,000 g for 10 min at 4°C. The supernatant was carefully removed and the pellet re-incubated for 30 min with 400 μl of methanol-acetic acid mix. Following the centrifugation, the supernatants were pooled. Extracts were then analyzed by LC-MS using an Acquity UPLC coupled to a XevoQtof (Waters, Massachusetts, USA). Analysis parameters were described in Jaulneau et al. (2010) [27].

###### Determination of Relative water content (RWC)

Tomato fully expanded leaves were excised and fresh weight (FW) was recorded every 30 minutes. The excised leaves were then allowed to float on deionised water for about 24 h and turgid weight (TW) was recorded. Leaves were dried at 80 °C for 24 h and dry weight (DW) recorded. Finally, RWC was calculated according to Smart and Bingham (1974) [28].

##### Quantitative expression assays

Three weeks WT and *ARF4*-as plants grown hydroponically in BD medium were subjected to either 150 mM of NaCl or 15% of PEG. Leaves and root samples were harvested after 2 hours and 24 hours for salt stress and after 48 hours for osmotic. RNA extraction, cDNA synthesis and real time PCR were performed as previously described in paragraph “ARF4 gene expression under salt and osmotic stress conditions”. The Gene-specific primers used are listed in Table S1.

## Results

### 3.1. An SlARF4 downregulated line shows altered anatomical, morphological and physiological parameters

We analyzed previously published *AUXIN RESPONSE FACTOR 4* antisense RNA post-transcriptionally silenced lines (*ARF4*-as) [14]. The *ARF4*-as plants exhibit a wide range of morphogenic phenotypes, namely delayed flowering, increased height and leaf curling as compared to their isogenic Micro-Tom (wild-type, WT) counterparts (Figure 1). Stem and root dry weight, on the other hand, showed a significant reduction in *ARF4*-as plants compared to WT, along with an insignificant increase in leaf area and specific leaf area (Figure 2).

**Figure 1.**
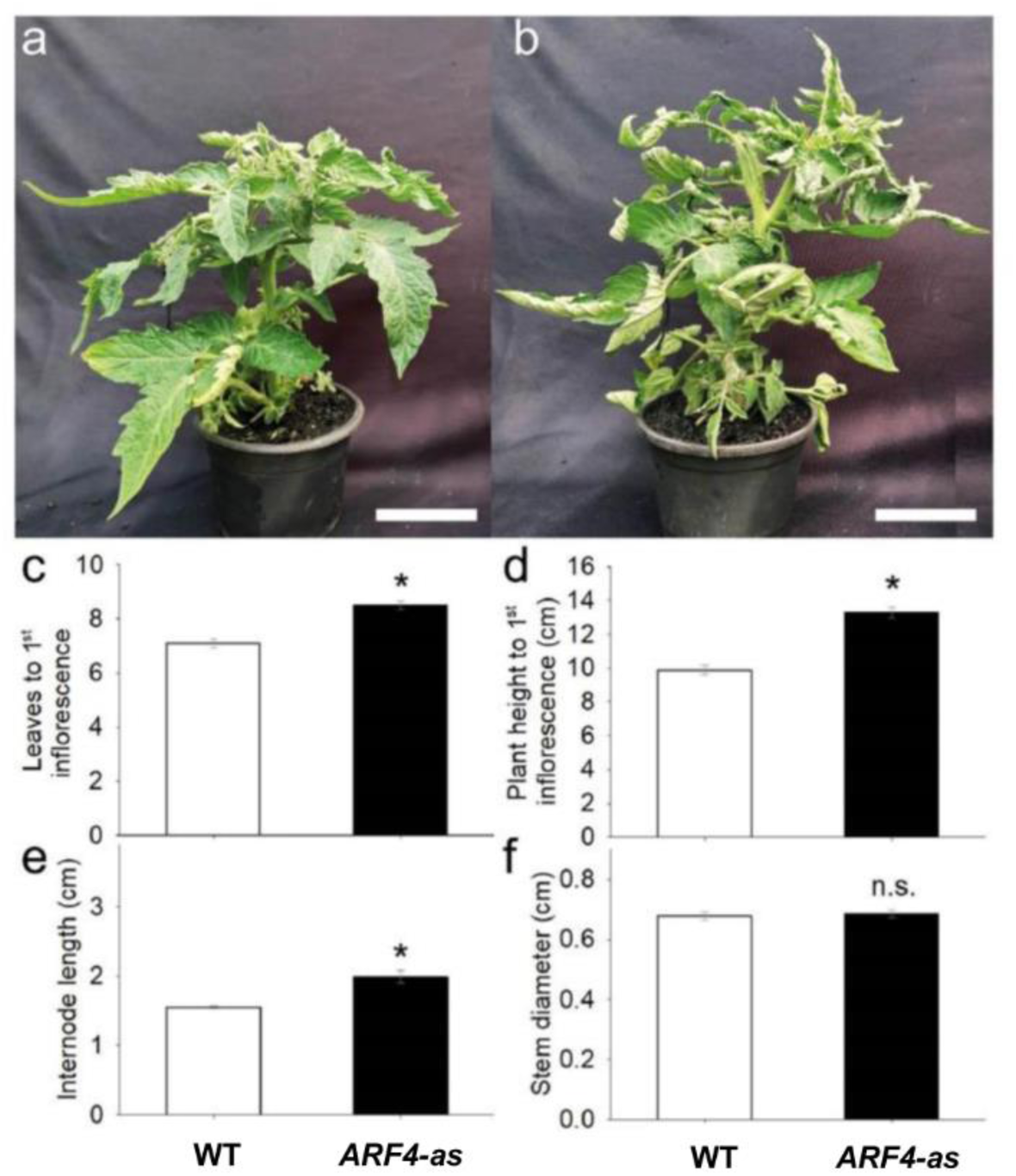
Phenotype of cv. Micro-Tom tomato plants (wild-type, WT) and isogenic *ARF4* antisense transgenic line (*ARF4-*as), 35 days after germination. (a) WT and (b) *ARF4*-as plants at the same stage of development, (c) number of leaves to first inflorescence, (d) plant height to first inflorescence, (e) plant internode length and (f) stem diameter. Bars are mean values (n=7) ± s.e.m. Asterisks indicate values that were determined by Student’s *t* test to be significantly different (P < 0.05) from WT.

**Figure 2.**
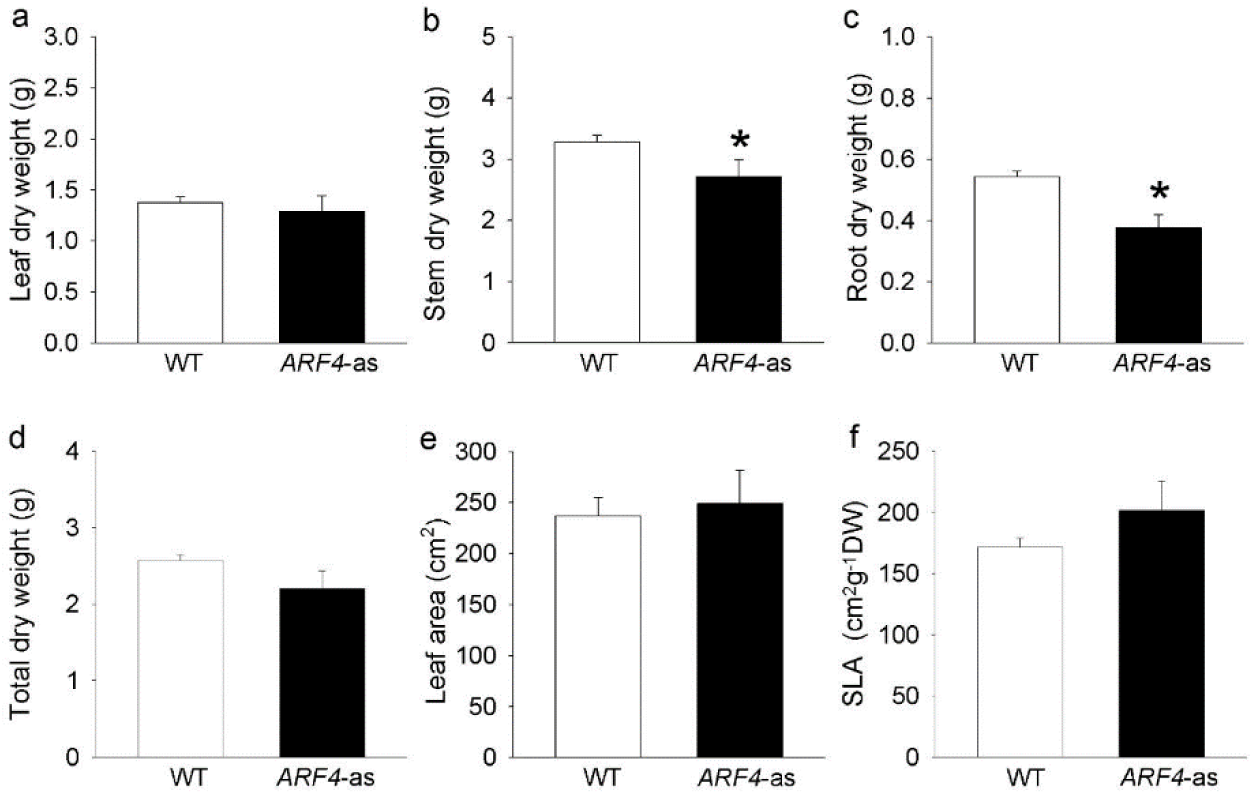
Dry weight and parameters related with leaf area for Micro-Tom (WT) and isogenic *ARF4* antisense transgenic line (*ARF4-*as), 45 days after germination. (a) leaf dry weight, (b) stem dry weight, (C) root dry weight, (d) total dry weight, (e) total leaf area and (f) specific leaf area (SLA). Values are means ± s.e.m (n=8). Asterisks indicate values that were determined by Student’s *t* test to be significantly different (P < 0.05) from the Micro-tom (WT).

*ARF4*-as plants analyzed in this work displayed an upward curling along the longitudinal axis of leaves (Figure 1b-3k). The extent of the changes detected in leaf morphology prompted us to check leaf ultrastructure in WT and *ARF4*-as plants. Leaf thickness was clearly reduced in *ARF4*-as plants as compared to WT (Figure 3a, b). In fact, the average of leaf blade thickness was 209 ± 8 µm in WT compared to 181 ± 7 µm in *ARF4*-as (Figure 3c). The palisade parenchyma (PP), spongy parenchyma (SP) and mesophyll tissue layers were thinner in *ARF4*-as plants than in WT plants (Figure 3d, 3e, 3j). The palisade:spongy mesophyll ratio was significantly higher in WT than in *ARF4*-as (Figure 3f). Meanwhile, the intracellular air spaces (IAS) were more conspicuous in *ARF4*-as compared to WT (Figure 3g), while no visible difference was observed in adaxial or abaxial epidermis between genotypes (Figure 3h-i).

**Figure 3.**
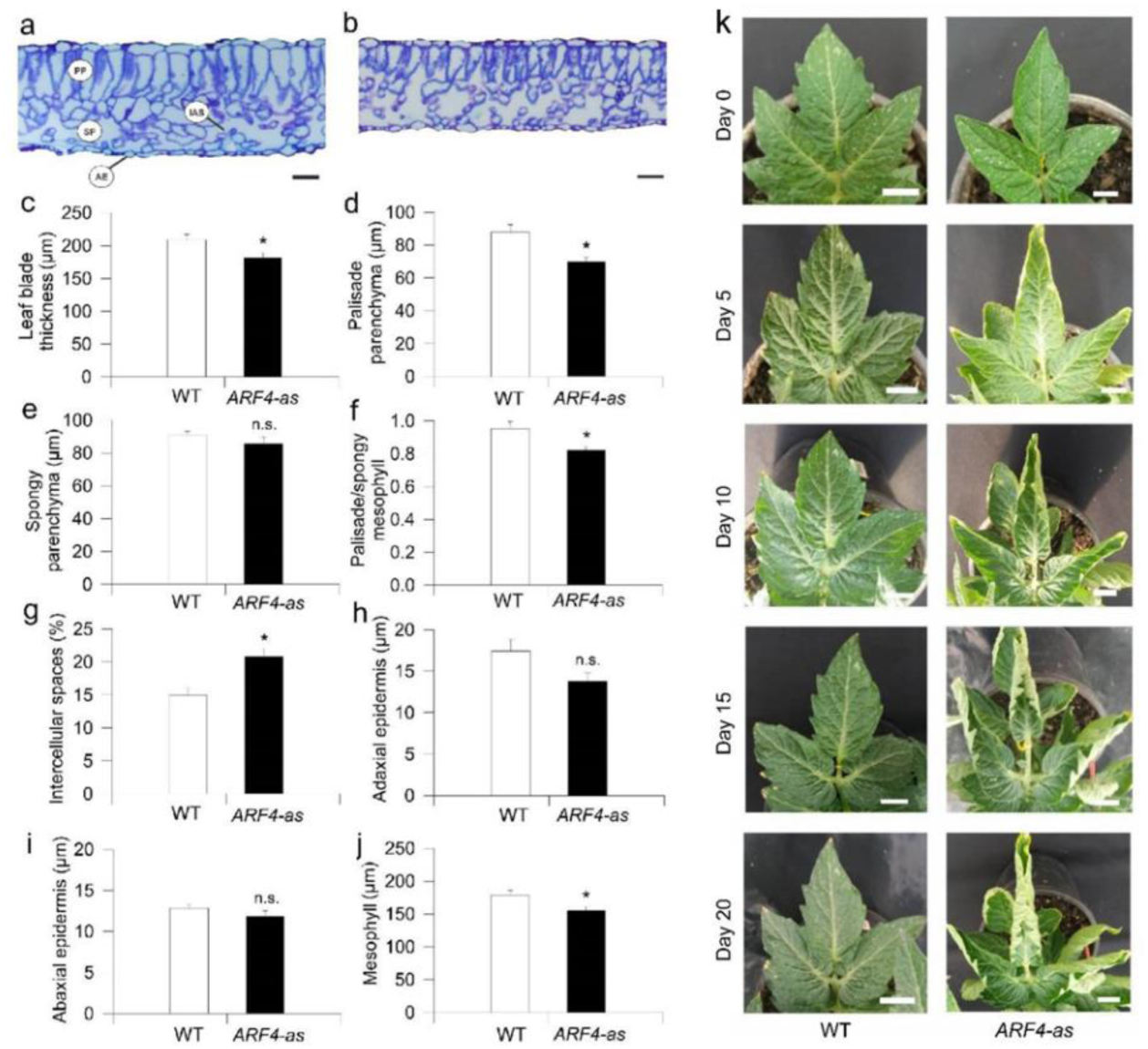
Leaf anatomy and morphology is altered in an *ARF4*-as transgenic line. Representative leaf cross-sections of (a) Micro-Tom (WT) and (b) *ARF4*-antisense line (*ARF4*-as). Scale bars= 20 µm. (c) Total thickness of the leaf blade, (d) thickness of palisade, (e) thickness of spongy and (f) ratio of palisade to spongy mesophyll, (g) proportion of intercellular air spaces, (h) adaxial and (i) abaxial and (j) mesophyll thickness in WT and *ARF4*-as plants and (k) Time series of a representative leaf illustrating blade curling in ARF4-as plants. Day 0 represents the day when the leaf was fully expanded. Values are means ± s.e.m (n=6).

Consistent with the presence of thinner leaves in the *ARF4*-as mutant plants, net (*A*) showed lower values along with a decreased stomatal conductance (*g*_s_) in *ARF4*-as plants while WT plants exhibited higher CO_2_ assimilation rate and *g*_s_ values (Figure 4a-b). It is well known that *A* can be limited by the slowest of two biochemical processes: (1) the maximum rate of ribulose 1,5-biphosphate (RuBP) carboxylase/oxygenase (Rubisco) catalyzed carboxylation (*V*_cmax_) and (2) regeneration of RuBP controlled by electron transport rate (*J*_max_) [29]. Consistently, *V*_cmax_ and *J*_max_ exhibited a significant decrease in *ARF4*-as plants (Table S2), indicating that photosynthesis is reduced due to the silencing of *SlARF4* (Figure 5a).

**Figure 4.**
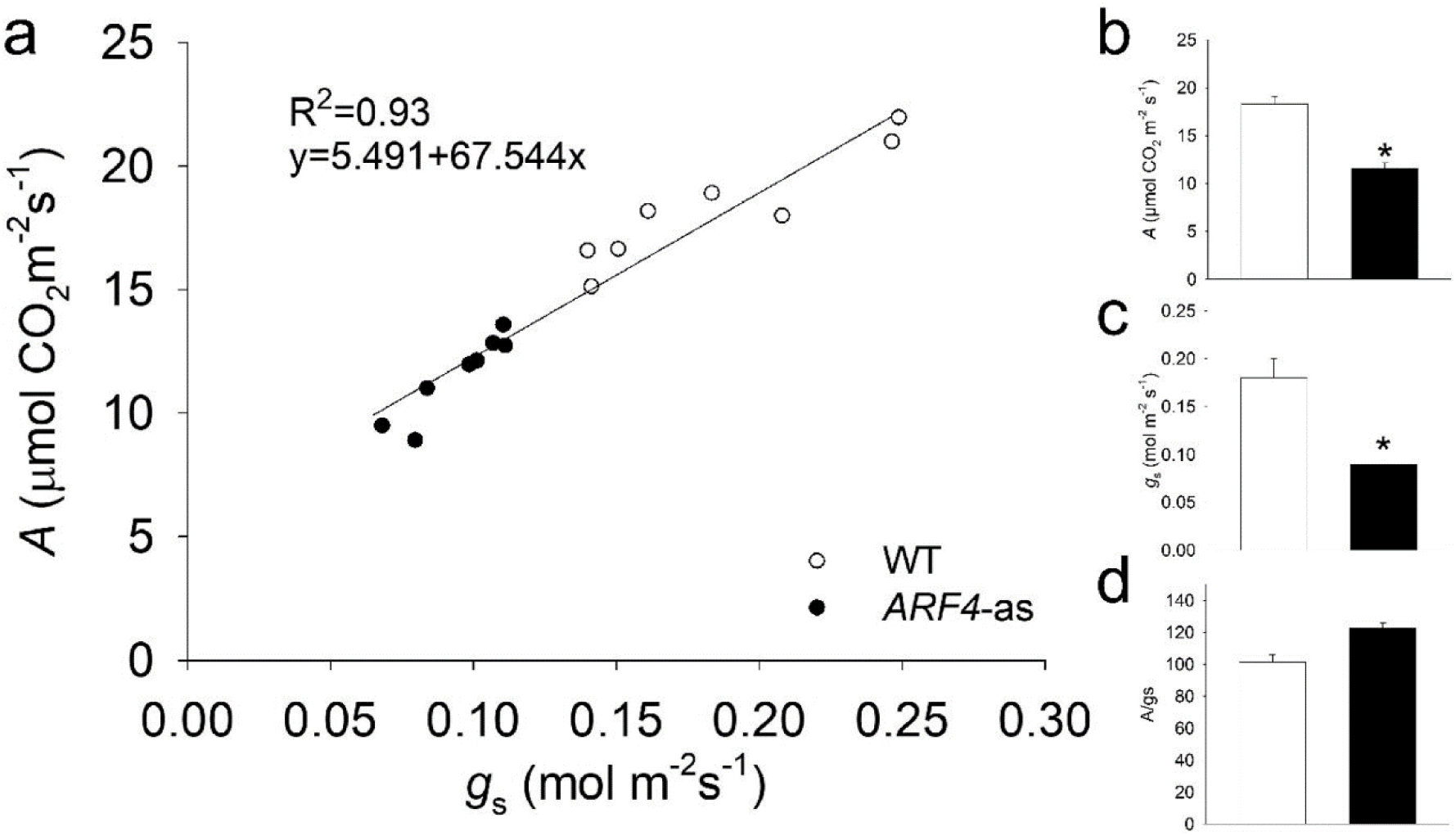
Net CO_2_ assimilation rate (A) and stomatal conductance (g_s_) in Micro-Tom (WT) and isogenic ARF4 antisense transgenic line (*ARF4*-as). (a) CO_2_ assimilation rate (A) as a function of stomatal conductance (gs), each point represents one measurement on an individual plant. Line fitted by linear regression. (b) net CO2 assimilation rate (A). (c) stomatal conductance (g_s_). (d) intrinsic water efficiency (A/g_s_). Values are means ± s.e.m (n=8). Asterisks indicate values that were determined by Student’s t test to be significantly different (P < 0.05) between genotypes.

**Figure 5.**
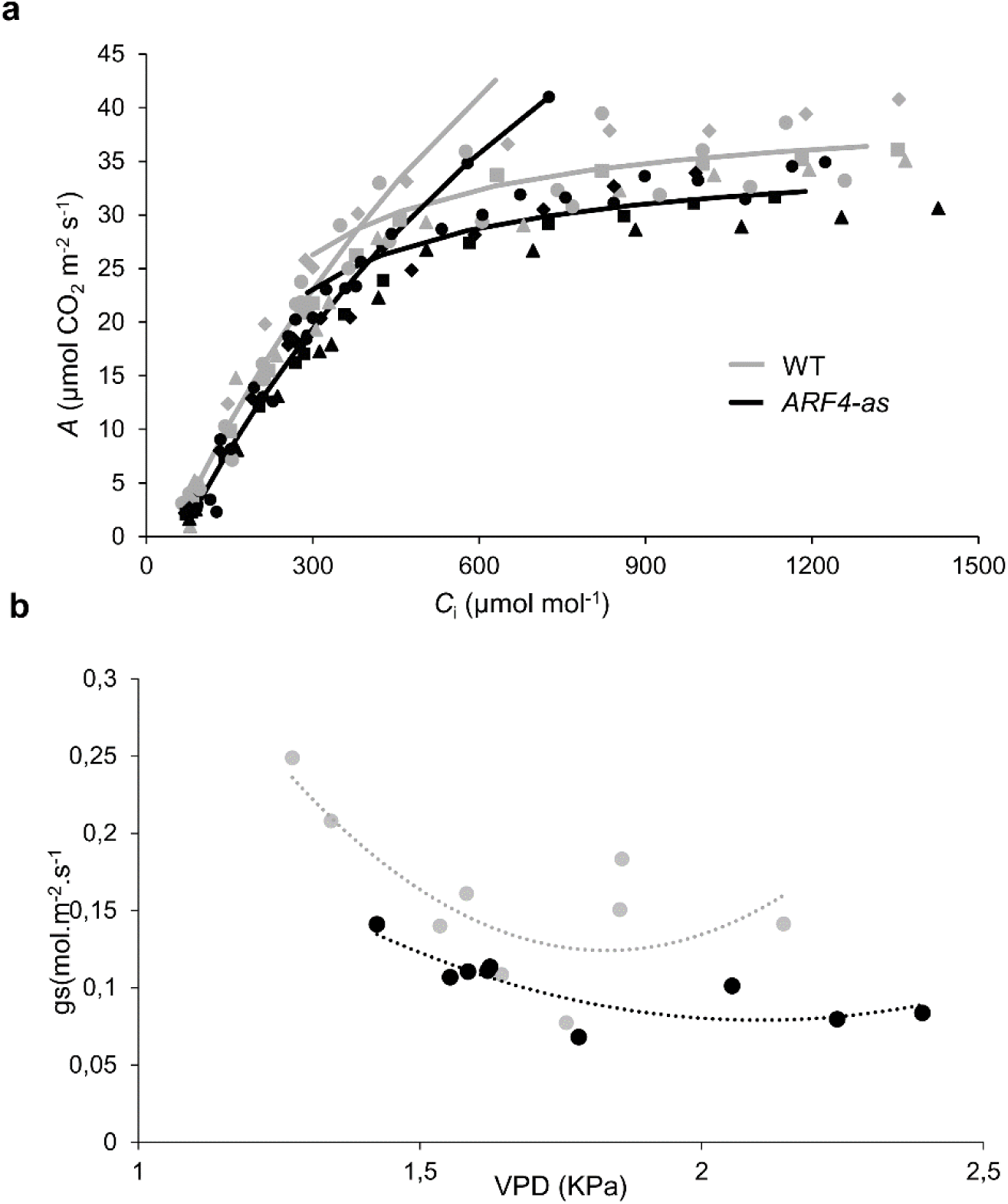
Photosynthetic assimilation rate and stomatal conductance in Micro-Tom (WT) and *ARF4*-antisense silencing line. (a) Net photosynthesis (*A*) curves in response to sub-stomatal (*C*_i_) CO_2_ concentration in WT and *ARF4-*as plants. Values are presented as means ± s.e.m. (n=5) obtained using the fully expanded fifth leaf. Two-branch curves: the biochemically based leaf photosynthesis model (Farquhar et al., 1980) was fitted to the data based on *C*_i_, values of *A*/*C*_i_ for five plants of WT (gray) and *ARF4-as* (black). (b) Response of stomatal conductance (*g*_s_) to changes in leaf-to-air vapor pressure deficit (VPD). Vapor pressure difference was varied by changing the humidity of air, keeping leaf temperature constant. Each point represents one measurement on an individual plant.

Stomatal conductance is one of the main parameters that affect photosynthesis under osmotic stress [30]. Because stomatal behavior is closely linked to CO_2_ uptake, it is considered to be one of the main reasons for reduced photosynthesis [31]. Stomatal conductance was significantly lower in *ARF4*-as as compared to WT (Figure 4c). The response of stomatal conductance (*g*_s_) to changing air vapor pressure deficit (*VPD*) was determined in *ARF4*-as and WT plants. At low *VPD*, WT plants exhibited high *g*_s_ and tended to reduce *g*_s_ as *VPD* increased from 1.25 kPa to 2 kPa and maintained a moderate *g*_s_ values at higher *VPD*s. Meanwhile, *ARF4*-as plants presented a relatively low *g*_s_ values at both low and high *VPD* compared to wild-type plants (Figure 5b). We verified a similarly impaired stomatal response in *ARF4*-as plants in response to step changes in irradiance and CO_2_ levels (Figure S1), suggesting that *ARF4* could be an important player in the regulation of stomatal movements. Furthermore, intrinsic water-use efficiency was notably higher in *ARF4*-as plants (Figure 4d) suggesting that the downregulation of *SlARF4* has the potential to improve the ration of carbon assimilation to transpirational water loss.

Given this potential impact on carbon assimilation, we next sought to examine whether *SlARF4* under-expression might affect plant agronomic parameters. Yield, number of fruits and fruit fresh weight were not altered between WT and *ARF4*-as plants, whereas fruit shape was altered in *ARF4*-as plants (Figure S2), as expected, given the well-known role of auxin in fruit development [11,32].

### 3.2. ARF4 has altered expression in response to salt and osmotic stresses

The expression pattern of *SlARF4* was analyzed by real-time PCR in leaves and roots exposed to salt or drought stress. *ARF4* gene expression was significantly regulated by salt and drought stresses. *SlARF4* expression was significantly repressed in leaves and roots after 2hours of salt treatment while its expression was significantly induced in leaves and roots after 24 hours (Figure 6a-b). During drought treatment, *SlARF4* gene was slightly induced after 48 hours of stress application in leaves (Figure 6c). In drought stressed roots, *SlARF4* was although significantly induced after 5 days of stress exposure (Figure 6d). Tomato lines harboring *pARF4*∷*GUS* constructs were generated and used to assay *in planta* analysis of *ARF4* expression under stress conditions. We observed strong expression in leaves, primary root and root tip after 24 hours of salt stress exposure (Figure 6e). After five days of osmotic treatment, GUS activity was detected in the root system and in different parts of the leaf (Figure 6f).

**Figure 6.**
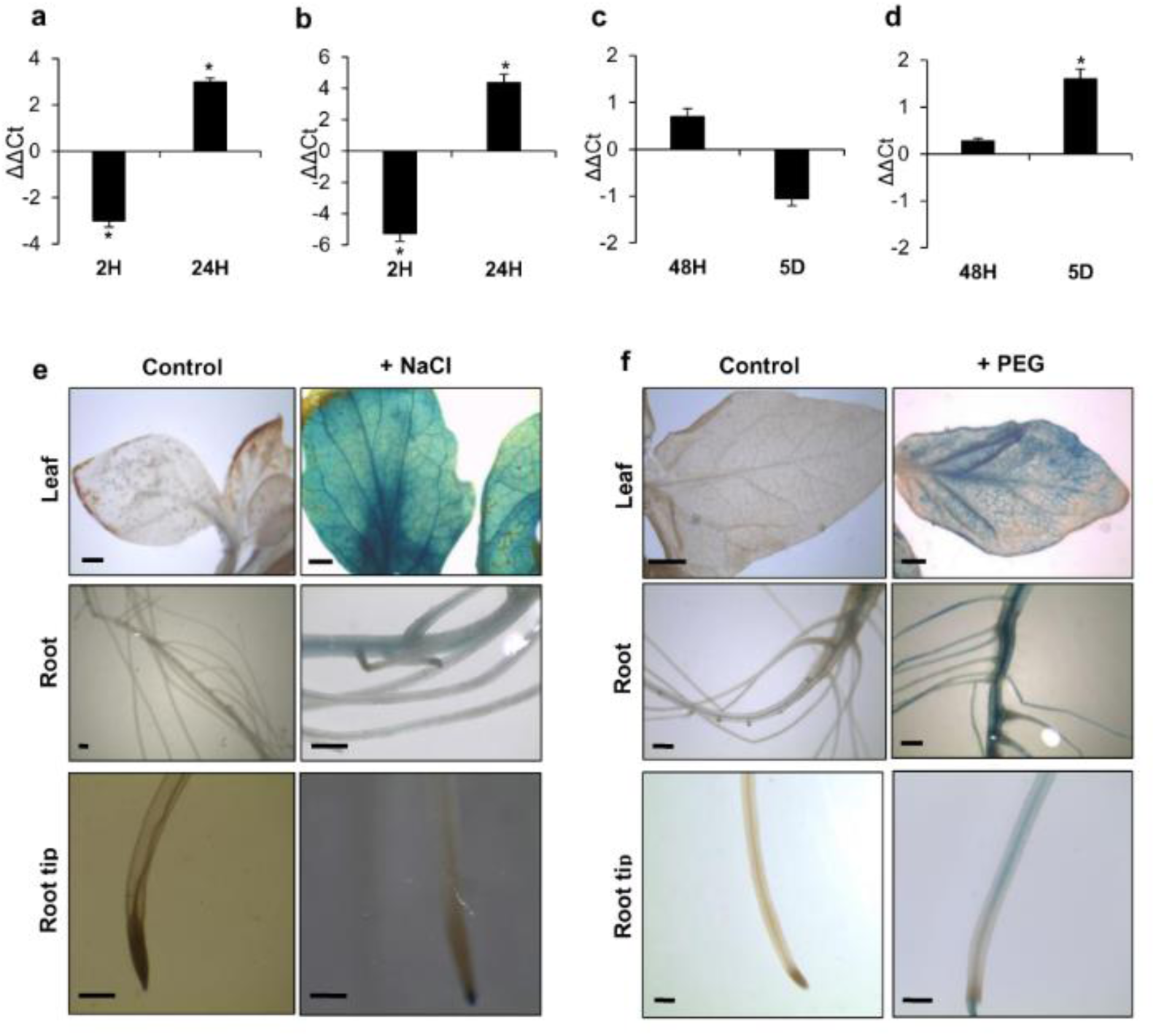
*SlARF4* expression in response to salt and drought stresses. Gene expression in leaves (a) and roots (b) of WT plants exposed to salt stress, in leaves (c) and roots (d) of WT plants exposed to osmotic stress. GUS activity in pARF4∷GUS tomato lines in salt (e) or osmotic (f) stress conditions. Bars scale (1mm). Stars (*) indicate the statistical significance (p<0,05) according to Student’s t-test.

### 3.3. ARF4 alters the plants response to salt and osmotic stress

#### 3.3.1. Shoot and root fresh weight are differentially altered in *ARF4* transgenic lines

Shoot fresh weight was investigated in WT and *ARF4-as* plants under salt and osmotic stress conditions. In the absence of stress, fresh weight was significantly higher in the *ARF4*-*as* as compared to WT (Figure 7a). Two levels of NaCl (100 mM and 150 mM) were applied, and both led to reductions in shoot fresh weight in both genotypes. In response to 150 mM NaCl, shoot fresh weight decreased by 60% in WT plants respectively and only by 30% in *ARF4*-as plants (Figure 7a). In roots, the reduction in the fresh weight was around 55% and 28% for WT and *ARF4*-as plants lines respectively (Figure 7c). Similarly, to salt stress, osmotic stress affected plant growth. Shoot fresh weight decreased by 66% for WT by 44% for *ARF4*-as plants when exposed to 15% of PEG 20000 (Figure 7b). Root fresh weight deceases significantly with the increase of PEG concentration. The decrease reached 57% in WT plants while the reduction was around 40% in *ARF4*-as in response to 15% of PEG20000 (Figure 7d).

**Figure 7.**
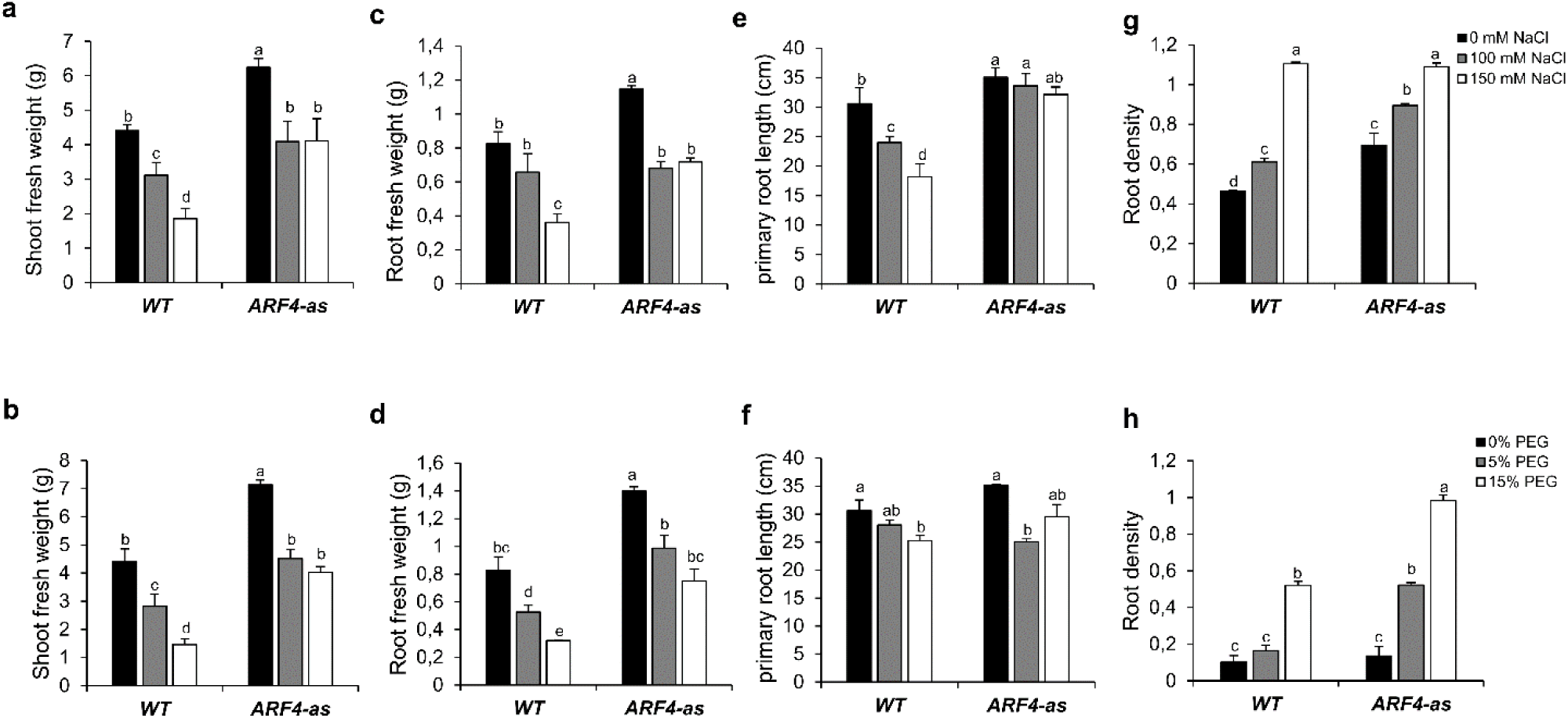
Growth parameters of tomato wildtype (WT) and *ARF4*-as in response to salt and osmotic stresses. (a) and (b) shoot fresh weight in salt and osmotic stress conditions respectively, (c) and (d) root fresh weight in salt and osmotic stress conditions respectively, (e) and (f) primary root length root in salt and osmotic stress conditions respectively, (g) and (h) root density in salt and osmotic stress conditions respectively. Salt and osmotic stresses were performed on three weeks tomato plants for two weeks by adding 100 mM of NaCl or 150 mM of NaCl for salt stress or 5% or 15% of PEG 20,000 for osmotic stress. Values are mean ± SD of three biological replicates. Bars with different letters indicate the statistical significance (p<0,05) according to Student Newman-Keuls test.

#### 3.3.2. Root development and density are less affected in *ARF4*-as plants

Root development was investigated in WT and *ARF4-as* plants in response to different concentrations of NaCl or PEG. Our results showed a significant reduction in primary root length in WT (by 40% in salt stress condition and by 17% in response to osmotic stress) while the decrease was nearby 8% and 12% in *ARF4*-as plants in response to salt and osmotic stress respectively (Figure 7e-f).

Root density was also investigated in the three lines under salt and osmotic stress conditions. Our results showed that root density significantly increased with the increase of NaCl or PEG concentrations in WT and in *ARF4*-as plants (Figure 7g-h). Root density increases by 57% in WT and 120% in *ARF4*-as., respectively. In response to osmotic stress, both plant lines showed a significant increase in root density. At 5% of PEG 20,000, root density increased by 50% and 130% in WT and *ARF4*-as plants respectively.

#### 3.3.3. *ARF4*-as plants are less affected by salt and osmotic stress

##### Photosynthesis is less affected in *ARF4*-as plants

Photosynthesis is among the primary processes to be affected by salinity and drought stress [33]. Our results showed that total chlorophyll content decreased significantly in WT plants exposed to different concentrations of NaCl while no significant changes in chlorophyll content was detected in *ARF4*-as plants (Figure S3). At 150 mM of NaCl, total chlorophyll content decreased only by 4% in the *ARF4*-as plants as compared to the control. Meanwhile, the diminution of total chlorophyll content was around 62.8% respectively in WT at 150 mM of NaCl compared to the control. The drastic decline in total chlorophyll content was also observed as a result of osmotic stress application on WT plants. The reduction in total chlorophyll content averaged the 37% in WT plants cultivated at 15% of PEG 20,000 respectively (Figure S3). Meanwhile, total chlorophyll content increased significantly*ARF4*-as in response to osmotic stress. In fact, the average of chlorophyll content was 1.2 times higher in *ARF4*-as at 15% PEG 20,000 than at 0% PEG 20,000.

##### Sugars are highly accumulated in *ARF4*-as plants in stress conditions

Soluble carbohydrates content was next investigated in *ARF4*-as and WT plant lines in response to different concentrations of NaCl or PEG. In response to salt stress, leaf soluble sugar content decreased by 57. 5% in WT plants and was increased by 146% in *ARF4*-as line exposed to 150 mM of NaCl for 2 weeks (Figure S3). In roots, soluble sugar content was also decreased in WT by 49. 4%, while an increase around the 191% in soluble sugar content was detected in *ARF4*-as plants. PEG induced osmotic stress imposed to plants significantly decreased soluble sugar contents at all the stress levels in WT and *ARF4*-as plants. Leaf soluble sugar content decreased by 42.8%and 18% respectively in WT and *ARF4*-as plants exposed to 15% of PEG 20,000. In roots, PEG application induced a significant decrease in soluble sugar content by 41.1% (at 15% of PEG 20,000) in WT plants respectively. Meanwhile, a significant increase in soluble carbohydrates content was although reported in *ARF4*-as plants and reached the 67% at 15% PEG 20,000 as compared to the control.

Carbohydrates produced in leaves during photosynthesis are transported throughout the plant by SUTs in order to support plant growth. The *SUT1* proteins are involved in the movement of sucrose into and out of the source and sink tissues and through the phloem via apoplastic pathways [34]. Investigating the expression of the sucrose transporter *SlSUT1* under salt and osmotic stresses had revealed that its expression was induced in WT and *ARF4*-as leaves and roots exposed to salt stress (Figure S4). These findings were also detected in response to osmotic stress. In osmotic stressed leaves, *SlSUT1* was significantly induced in WT and *ARF4*-as plants. In roots, the expression of *SlSUT1* was also up-regulated significantly.

##### *ARF4*-as plants showed lower stomatal conductance

Stomata play an imminent role in plant response to environmental changes as they control both water losses and CO_2_ uptake [35]. Stomatal conductance was estimated in WT and *ARF4*-as plants in salt or osmotic stress (Figure 8a-b). Under normal conditions, *ARF4*-as plants exhibited low stomatal conductance as compared to WT plants. Salt and osmotic stress application induces a significant decrease in stomatal conductance in WT. In fact, the reduction in stomatal conductance observed in the WT was around 32% and 46% in response to salt and osmotic stress respectively. Meanwhile, no significant changes in stomatal conductance were detected in *ARF4*-as plants that still maintained the same values of stomatal conductance observed in normal conditions.

**Figure 8.**
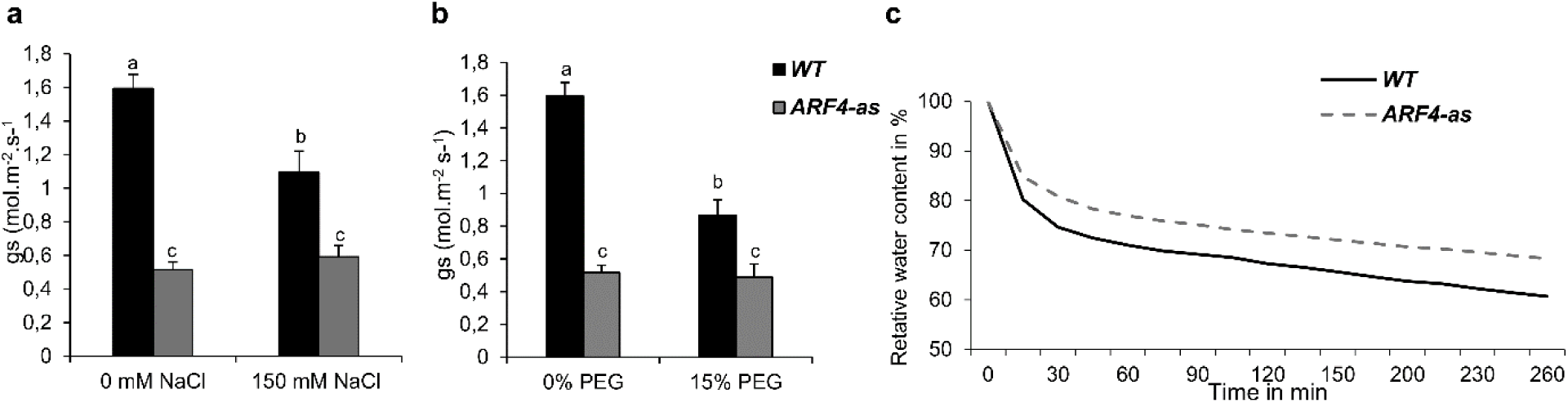
Stomatal conductance and relative water content in WT and *ARF4*-as plants. (a) and (b) stomatal conductance in salt and osmotic stress conditions, (c) relative water content in response to dehydration. Bars with different letters indicate the statistical significance (p<0,05) according to Student Newman-Keuls test.

Stomatal closure can be trigged by various internal and external factors. Schultz (2003) have shown that the decline in leaf water potential can trigger stomatal closure under stressful conditions [36]. Relative water content (RWC) was investigated in WT and *ARF4*-as plants. A remarkable decrease in RWC was observed in both lines (Figure 8c). Up to 25% and 15% of water losses was recorded during the first 30 minutes of dehydration in WT and *ARF4*-as plants respectively. RWC reached 72% in WT and after 4 hours of dehydration while a 50% decrease of water losses was found in *ARF4*-as plants.

##### *ARF4*-as plants exhibited a high ABA content

As ABA is the key hormone for regulating stomatal aperture and thus RWC [37], endogenous ABA content of 5 weeks *ARF4*-as and WT plants in normal or under salt and osmotic conditions were then measured in leaf tissues. Our results showed that the ABA content in *ARF4*-as increased by 120% and 101% under salt and osmotic stress conditions respectively whereas in WT plants increased by only 20% in salt conditions and decreased by 26% in PEG treatment (Table 1).

**Table 1:**
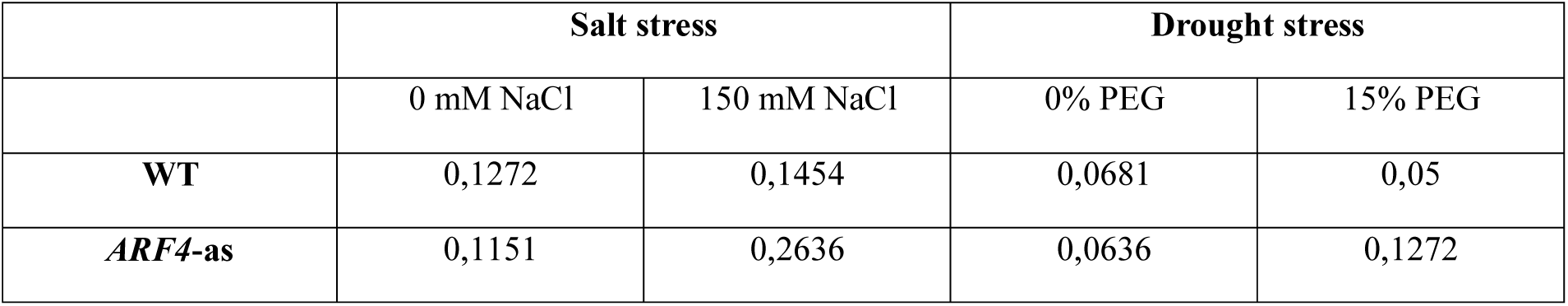
ABA content (expressed in µmol/g of FW) in WT and *ARF4*-as leaves in response to salt or osmotic stress conditions. Values presented are mean ± SD of three biological replicates (except for data related to *ARF4*-as plants in osmotic stress conditions, where only two biological replicates were done).

Endogenous ABA content is related to the balance between its biosynthesis and its degradation. In tomato, *SlNCED1* and *SlNCED2* encoding 9-cis-epoxycarotenoid dioxygenase are the primary genes responsible for ABA biosynthesis, whereas *SlCYP707A1, SlCYP707A2, SlCYP707A3* and *SlCYP707A4* encoding ABA 8′-hydroxylase are main genes for ABA catabolism [38]. Investigating the expression of these genes in WT and the *ARF4*-as plants had revealed that their expressions were significantly regulated by salt or osmotic stress application (Figure 9). Indeed, *SlNCED1* expression was significantly induced in *ARF4*-as leaves after 2 hours and 24 hours of salt stress exposure while no significant changes was observed in roots. In WT, the expression of *SlNCED1* was significantly up regulated in leaves and roots after 24 hours of salt treatment. In response to osmotic stress, the expression of *SlNCED1* was significantly up-regulated in *ARF4*-as roots while no significant change in *SlNCED1* expression was detected in WT plants. *SlNCED2* expression was significantly induced in the *ARF4*-as leaves after 2 hours and 24 hours of salt application and in WT after 24 hours. A significant increase in the expression of *SlNCED2* was detected in *ARF4*-as plants in response to osmotic stress.

**Figure 9.**
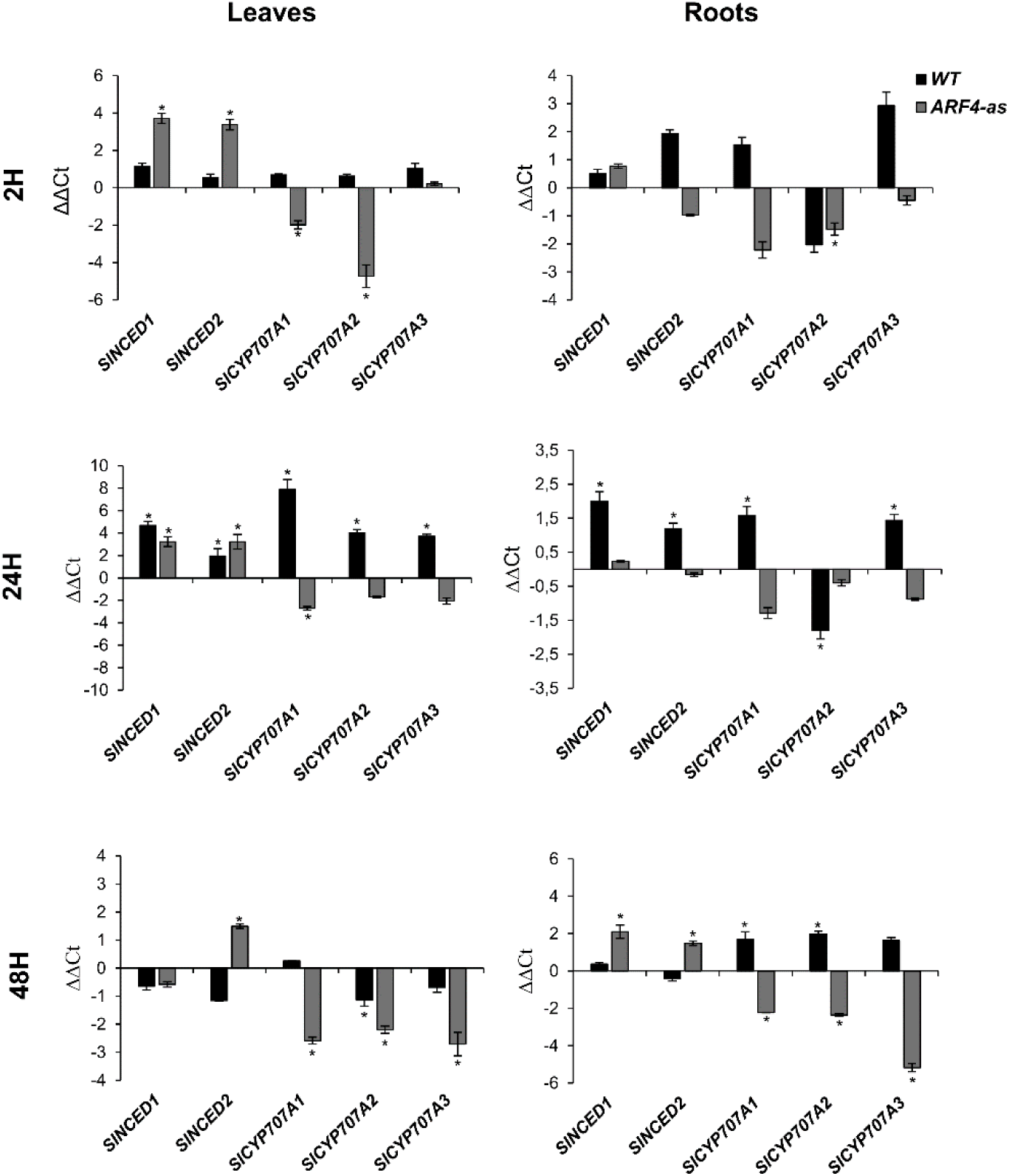
Expression of *SlNCED1, SlNCED2, SlCYP707A1, SlCYP707A2* and *SlCYP707A3* genes in WT and *ARF4*-as leaves and roots after 2H and 24H of salt stress and 48H of osmotic stress application. ΔΔCt refers to fold differences in gene expression relative to untreated plants. Values presented are mean ± SD of three biological replicates. Asterisks (*) indicate the statistical significance (p<0,05) according to Student’s t-test.

The expression pattern of *SlCYP707A1, SlCYP707A2* and *SlCYP707A3*, key ABA degradation genes was also assessed in *ARF4*-as and WT plants under salt and osmotic stress conditions (Figure 9). Quantitative RT-PCR results showed that the expression of the ABA metabolism genes was up-regulated in WT leaves after 2 hours and 24 hours of salt stress exposure. In roots, the expression of *SlCYP707A1* and *SlCYP707A3* was up regulated in WT plants in response to salt stress while a significant repression was reported in the expression of *SlCYP707A2*. Although, the expression of the ABA metabolism genes expression was repressed in the *ARF4*-as salt stressed roots. In response to osmotic stress, the expression of *CYP707A1, CYP707A2* and *CYP707A3* was repressed in *ARF4*-as plants while their expression was up regulated in WT roots and downregulated in leaves.

##### Antioxidant genes expression is altered in response to salinity and osmotic stress

###### *Cat1* (Catalase) expression in response to salt and osmotic stress

Catalases are tetrameric heme-containing enzymes with the potential to directly dismutate H_2_O_2_ into H_2_O and O_2_ and essential for ROS detoxification [39]. In angiosperms, catalase is encoded by three genes known as *Cat1, Cat2* and *Cat3* [40]. We investigated *Cat1* expression in WT and *ARF4*-as plants exposed to salt and osmotic stresses. Our results showed that *Cat1* expression was induced in *ARF4*-as plants exposed to salt stress. A high up regulation in *Cat1* expression was although detected in WT plant roots in response to salt stress (Figure S5). In osmotic stressed leaves, *Cat1* expression was significantly induced in WT while a remarkable but not significant repression was observed in *ARF4*-as plants. Meanwhile, *Cat1* gene expression was highly up regulated or repressed in WT and *ARF4*-as plant roots respectively.

###### *SOD* (Superoxide dismutase) expression in response salt and osmotic stress

Superoxide dismutase (SOD) is an enzyme that belongs to the family of metalloenzymes ubiquitous in all aerobic organisms. It is one of the first defense against ROS induced damages by catalyzing the removal of O_2_^-^ by dismutating it into O_2_ and H_2_O_2_ [39,40]. Based on the metal ion it binds, SOD are classified into three isozymes; Mn-SOD (in mitochondria), Fe-SOD (in chloroplasts) and Cu/ZnSOD (in cytosol, peroxisomes and chloroplasts) [39]. Investigating the expression of *Cu/ZnSOD* had revealed that its expression was remarkably repressed in WT plants subjected to salt and osmotic stresses. Meanwhile, *Cu/ZnSOD* gene expression was induced in *ARF4*-as leaves in response to salt and osmotic stresses while its expression was repressed in the roots (Figure S5).

###### *mdhar* (Monodehydroxyascorbate reductase) expression in response to salt and osmotic stress

Monodehydroxyascorbate reductase (MDHAR) is another enzymatic antioxidant indirectly involved in the ROS scavenging through regenerating Ascorbic acid, indispensable for the dismutation of H_2_O_2_ by APX enzyme [39]. Investigating the expression pattern of *mdhar* gene in salt stress conditions had revealed a significant up regulation in *ARF4*-as leaves and roots while its expression was repressed in WT plants (Figure S5). In response to osmotic stress, the expression of *mdhar* was significantly up regulated and repressed respectively in WT. In roots, *mdhar* is significantly up regulated in *ARF4*-as plant line.

### 3.4. SlARF4-crispr mutant exhibited similar alteration in growth and stomatal functions observed in ARF4-as plants

*SlARF4* mutants were also generated using the revolutionary CRISPR-Cas9 technology. For that, the two target sequences were located in the *SlARF4* DBD domain. The CRISPR transformants from T1 generation of Line 108 were genotyped (by PCR and DNA sequencing). Based on PCR results, plants #5, #6 and #7 yielded a DNA fragment with a 49bp DNA deletion. The *Cas9* transgene was segregated out in plant #5 while still bearing the desired mutation of *SlARF4* gene. The presence of *Cas9* gene in *arf4-cr* mutants was confirmed in two of the three randomly analyzed plants (Figure 10a). DNA sequencing of *ARF4*-PCR product from plant #5 and #6 revealed that plant #5 harbored a single fragment type containing the expected 49bp DNA deletion. However, only four out of 10 PCR clones from plant #6 hold the 49bp deletion in the DBD domain while the remaining 6 PCR clones showed although a small deletion in both target regions (Figure 10b). At the morphological level, the T1 generation plants showed dramatic up curling phenotype, similar to the one observed in *ARF4-*as plants (Figure 10c, d).

**Figure 10.**
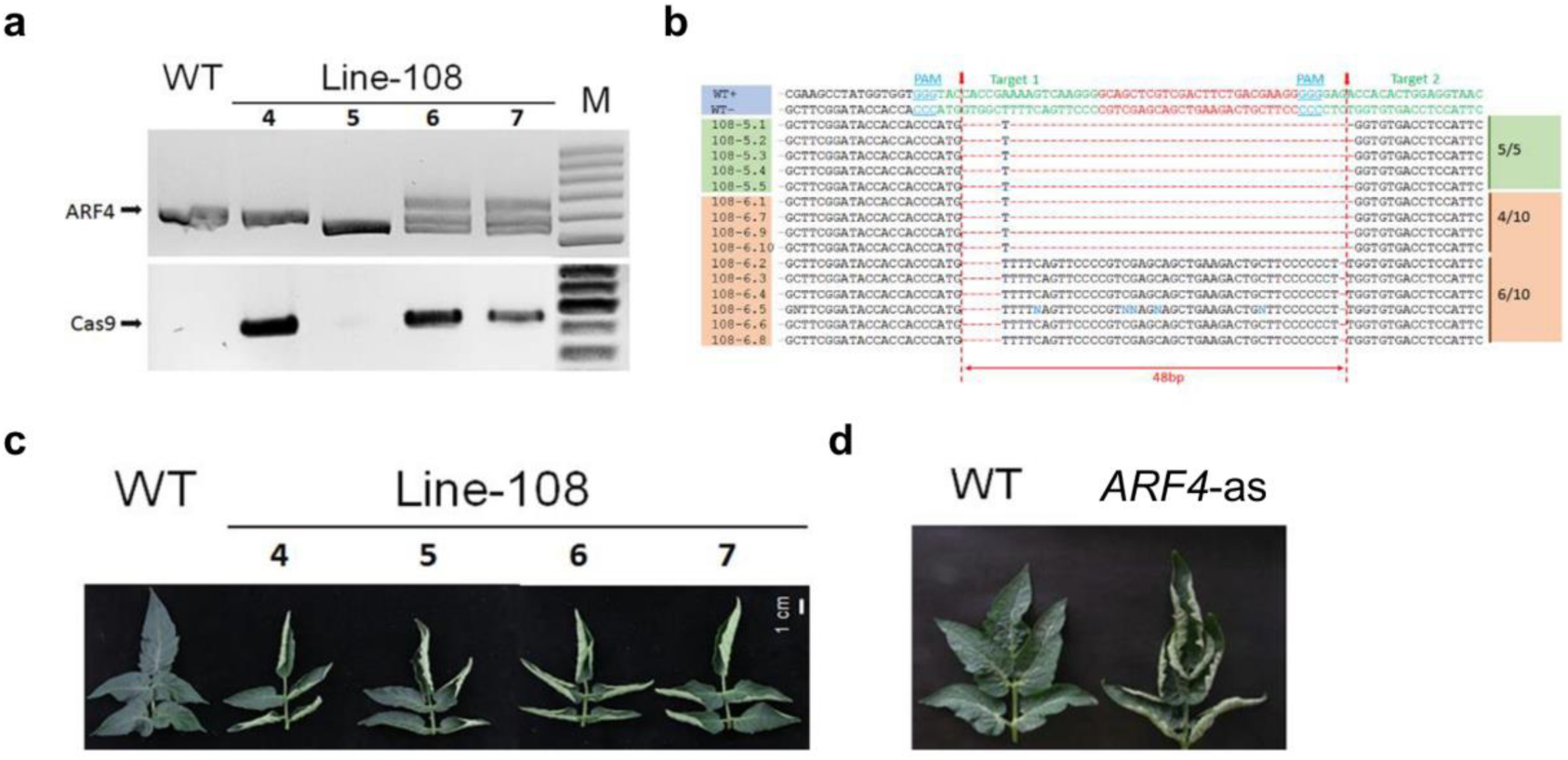
CRISPR-Cas9 mediated gene editing in tomato Micro-Tom. (a) PCR genotyping of plants at the T1 generation of line-108. Deletion mutations of *SlARF4* were found in plants #4, #5, #6 and #7. Among these, the T-DNA insertion (CRISPR-Cas9 transgene) was segregated out in plant #5 while still bearing the desired mutation in the *SlARF4* gene. (b) Sequencing data of *ARF4*-PCR products from #5 and #6 plants. The PCR products from plant #5 yielded a single fragment type containing the expected 49bp DNA deletion, whereas in the case of plant #6, only 4 out of 10 PCR clones contain the desired mutation, the remaining 6 PCR clones exhibited a small deletion in both target regions. Red dashed indicated the expected cleavage sites for CRISPR-Cas9. (c) Leaf phenotype of CRISPR-Cas9 generated *ARF4* mutants. All four plants showed dramatic up curling leaves, similar to the phenotype observed in *ARF4*-as plants (d).

Besides the up-curling leaf phenotype, *arf4-cr* mutant (#5) showed a significant decrease in leaf, root and stem dry weight as compared to WT siblings (Figure 11a, b, c). Leaf area was also significantly lower in *SlARF4*-Crispr plants (Figure 11 d). Moreover, as expected, net CO_2_ assimilation rate (A) and stomatal conductance (g_s_) was significantly lower in *arf4-cr* plants than in WT while water use efficiency was significantly higher in *SlARF4*-Crispr mutant (Figure 12). Regarding the agronomical traits, plant productivity and quality were not affected by *SlARF4* silencing by CRISPR technology. Yield and fruit fresh weight of *arf4-cr* plants were slightly but not significantly higher as compared to WT while equatorial and polar diameter of *SlARF4*-Crispr fruits were significantly higher (Figure S6). Most of these growth and stomatal alterations detected in *arf4-cr* plants was also found in *ARF4*-as plants (Figure 2, 4, S3).

**Figure 11.**
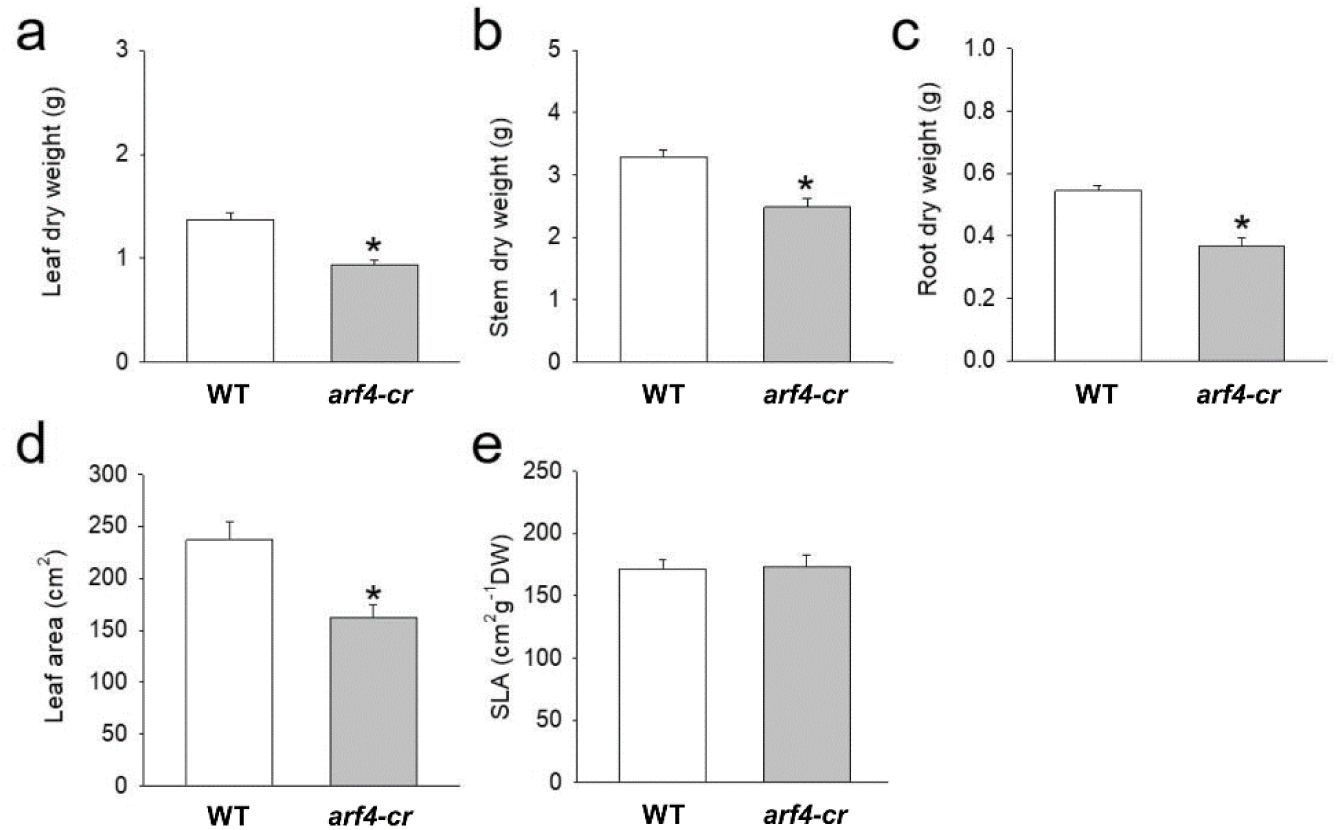
Dry weight and parameters related with leaf area for Micro-Tom (WT) and *arf4-cr*, 45 days after germination. (a) leaf dry weight, (b) stem dry weight, (c) root dry weight, (d) leaf area and (e) Specific leaf area (SLA). Values are means ± s.e.m (n=8). Asterisks indicate values that were determined by Student’s t test to be significantly different (P < 0.05) from the Micro-tom (WT).

**Figure 12.**
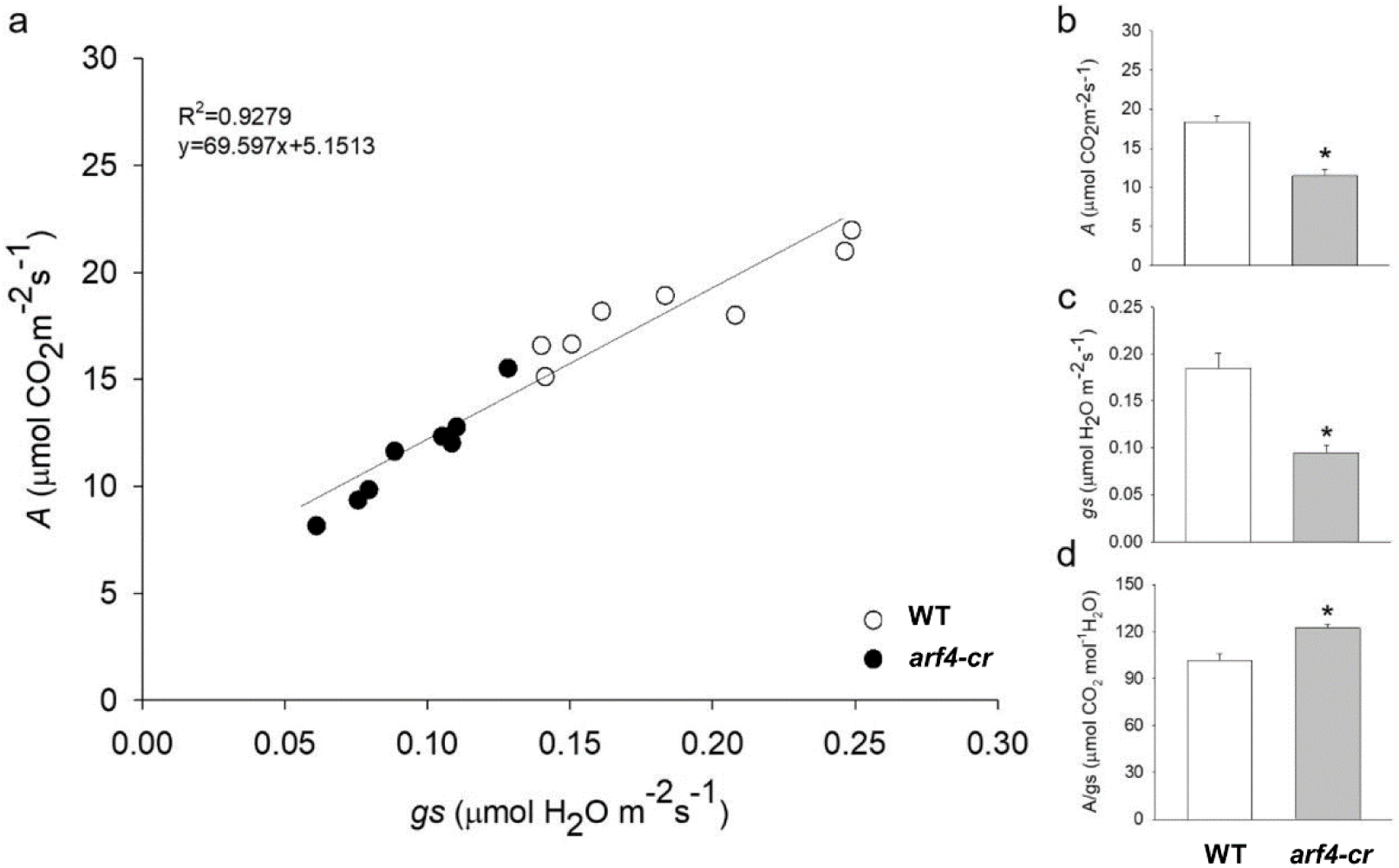
Relationship between net CO_2_ assimilation rate (*A*) and stomatal conductance (*g*_s_) for Micro-Tom (WT) and *arf4-cr*. (a) CO_2_ assimilation rate (*A*) as a function of stomatal conductance (*g*_s_), (b) net CO_2_ assimilation rate (*A*), (c) stomatal conductance (*g*_s_), (d) intrinsic water efficiency (*A*/*g*_s_). Values are means ± s.e.m (n=8). Asterisks indicate values that were determined by Student’s t test to be significantly different (P < 0.05) from the wild type (WT).

## 4. Discussion

In the face of a global scarcity of water resources and the increased water and soil salinization, abiotic stresses present major challenges in sustaining crop yield. Plant tolerance to these stresses relays on the implementation of molecular mechanisms for cellular adjustments, including signal perception and transduction cascades, transcriptional networks and adaptive metabolic pathways. Plant hormones such as ABA, ethylene and SA play an important role in mediating plant responses to stresses. Auxin, the key regulator of many aspects of plant growth and development, was identified as a factor in plant response to biotic and abiotic stress. ARFs play crucial role in auxin signaling. *ARF4* is involved in the regulation of several processes such as lateral root development, fruit quality and development [13,14,41]. Here, we demonstrated that the under expression of *ARF4* confers plants tolerance to salt and osmotic stress. Interestingly, even under stress conditions, *ARF4*-as plants show clearly (i) a proper plant development, (ii) a high chlorophyll content along with a significant accumulation of sugars, (iii) an increase in ABA content and a higher water-use efficiency, (iv) a significant upregulation of antioxidants and thus (iv) an enhanced stress tolerance compared to WT plants. These multiple fitness benefits obtained by under expressing *ARF4* in plants constitute a desired characteristic for transgenic crop plants nowadays.

Successful growth of plants relies on the plasticity in leaf anatomical characteristics, which enables plants to cope with diverse stress environments [42]. The transcriptional downregulation of *ARF4* expression leads to severe leaf curling along the longitudinal axis which was consistent with those observed in DR12-ASL lines generated in the Kemer cultivar by Jones *et al*. (2002) and those obtained in Micro Tom by Sagar *et al*. (2013) [14,43]. Leaf rolling is a one of the common adaptation to water stress [44]. It has been reported that greater leaf rolling may be an important indicator linked to osmotic tolerance and may have a positive impact on crop yield under water stress conditions [45]. Plants having leaf rolling mechanism exhibit a resistance to osmotic and high temperature and had higher water-use efficiency. This was explained by the fact that leaf rolling reduced transpiration rate [44].

Plants’ tolerance to abiotic stress depends also on their ability to adjust the relationship between water, transpiration, photosynthesis and water use efficiency through stomatal changes in order to maximize CO_2_ assimilation [46]. In fact, the decrease in stomatal conductance can enhance plant tolerance to osmotic stress of many plant species such as chickpea and rice [47,48]. *ARF4*-as plants showed a decrease in stomatal conductance coupled with an increase in water-use efficiency, suggesting that the downregulation of *ARF4* can lead to a better tolerance to abiotic stresses. Thus, the combination of a marked leaf curling phenotype associated with decreased *g*_s_ and high water-use efficiency in *ARF4*-as plants suggests that these plants might tolerate better salinity and water deficit.

The auxin response has emerged recently as an active actor in plant response to abiotic stresses [49]. Extensive research has shown that various environmental signals are associated with changes in auxin homeostasis, redistribution, and signaling [50]. Accumulating evidence indicates that auxin plays a role in plant responses to abiotic stresses through complex metabolic and signaling networks. Auxin coordinates plant development essentially through two transcriptional regulators Aux/IAA and ARFs [51]. Genome-wide expression analyses have suggested that the expression of numerous *ARF* genes change when plants respond to abiotic stresses in many plant species [6,8,52,53]. In tomato, expression profiling of the *ARF* family under abiotic stress conditions showed that most of the tomato *ARF*s were responsive and some of them were significantly regulated [16]. Among them, *SlARF4* expression was significantly induced in tomato plants in response to salt (24 hours) or drought (five days). Additionally, *SlARF4* regulatory region was enriched in *cis*-acting elements specific to salinity and water deficit response [16]. In this work, *ARF4* expression assessed both *in vitro* and *in planta* was affected by either salt, drought or osmotic treatment, which suggest that this gene is involved in tomato response to those stresses.

Plant growth and development are heavily constrained by salinity and water deficit [54]. In the current study, plant growth evaluated through fresh weight was significantly affected in WT and *ARF4*-as transgenic plants in response to salt or osmotic stress. The reduction in fresh weight was reported in many plant species which includes barley, cabbage and sorghum in response to salinity or water deficit [55–57]. The decrease in seedling growth (evaluated through the measurement of fresh weight) is due to restricted cell division and enlargement, as salt and drought stress directly reduces growth by decreasing cell division and elongation [58]. However, the reduction in fresh weight was less pronounced in the *ARF4*-as plants which suggest that *ARF4*-as plants might tolerate better salt and osmotic stress than the other plant lines.

The root system is the first organ to encounter salinity and drought stress. Root development can severely be affected by environmental stresses [59]. In *Arabidopsis*, root length and development was significantly reduced in response to salinity [60]. In this study, we had noticed a significant decrease in primary root length in WT and in *ARF4-as* plants in response to salt or osmotic stress conditions as compared with normal conditions. However, less reduction was reported in *ARF4*-as plants. In general, deeper rooting has been shown to be beneficial for plant production and survival as it increases water uptake which confer the advantage to support plant growth during adverse conditions [61]. Numerous studies have linked plant stress tolerance with the increase in root length and density for several plant species such as barley, sunflower, wheat, rice and cotton [62]. *ARF4*-as plants showed less reduction in primary root length coupled with an increase in root density. *ARF4* was found to be implicated in defining root architecture in *Arabidopsis* under optimal growth conditions through the control of lateral root emergence [41] which suggest that this gene might play an important role in root architecture in stress conditions and thus contribute to the improvement tomato tolerance to salinity and osmotic stress.

Plants’ photosynthesis activity is known to be affected by abiotic stress [63]. Stress-induced decrease in chlorophyll content have been reported in several plant species including tomato [64]. Zarafshar et al. (2014) explained this decrease as the result of pigment photo-oxidation and chlorophyll degradation [65]. *ARF4*-as plants showed although a slight decrease in total chlorophyll content in response to salt stress and a slight increase in osmotic stress conditions. Tomato tolerant genotypes showed less losses in photosynthetic pigments [66]. The lack of changes in chlorophyll content in the *ARF4*-as plants shows the capacity to preserve the photosynthetic apparatus and thus indicates their better tolerance to salinity and osmotic stress.

Metabolic activities in plant cells are very complex, and various biochemical pathways are interconnected with each other, working in coherence towards cellular homeostasis [67]. High concentration of osmolytes including sugars help plants to tolerate abiotic stresses by improving their ability to preserve osmotic balance within the cells [68]. To this purpose, we studied sugars accumulation to correlate their levels with the presence or the absence of *SlARF4* gene. We found that soluble sugars content (Fig. S3) increased more in *ARF4*-as transgenic plants than in WT in response to salt or osmotic stresses. Besides its accumulation under stress conditions, sugars can be transported to different organs of the plant. *SUT1*; gene encoding for a sucrose transporter, was found to be upregulated in *ARF4*-as plants in response to salt or osmotic stress. Previous studies had reported an increase in *SUT1* transcript in sugarcane leaves after 24 hours of osmotic treatment with PEG [69]. Thus, its upregulation in *ARF4*-as might improve tolerance to osmotic and salinity through the stimulation of sucrose transport required for the osmoregulation and for the cellular energy demands during salt or osmotic stresses.

Stomata are known for their role in the regulation of gas exchange and water loss by transpiration [70]. Their opening and closing is affected by environmental and internal parameters that maintain the water balance and functioning of complex signal transduction pathways [71]. Salt, drought and osmotic tolerance was correlated with a decline in stomatal conductance in many plant species [47,48]. In this work, we found that the downregulation of *SlARF4* resulted in reduced stomatal conductance under salt and osmotic stresses along with a high RWC, which suggests the involvement of *ARF4* in the regulation of stomatal closure in order to prevent water loss through transpiration to cope with salt or drought conditions.

Stomatal closure is a key ABA-mediated process for coping with water deficits. Stomatal closure can be trigged by the subsequent accumulation of ABA [72]. ABA-insensitive mutants (i.e., *abi1* and *abi2*) are very susceptible to drought mainly due to the impairment of their stomatal apertures ^79^. Thus, the lowest stomatal conductance observed in *ARF4*-as plants may be associated with the accumulation of relatively more ABA in *ARF4*-as stressed leaves and suggests that *ARF4*-as plants can tolerate better salt and osmotic stresses than WT plants.

ABA content is closely defined by the balance between its biosynthesis and biodegradation. In tomato, ABA biosynthesis is governed by the activity of *SlNCED1* and *SlNCED2*, whereas *SlCYP707A1, SlCYP707A2, SlCYP707A3* and *SlCYP707A4* encoding ABA 8′-hydroxylase are main genes for ABA catabolism [38]. Water stress is known to induce the expression of *NCED* genes in tomato [75]. Moreover, transgenic plants overexpressing the *NCED* gene accumulated large amounts of ABA and were more resistant to drought stress [76]. Regarding ABA biodegradation genes, *Arabidopsis* knock-out mutant *cyp707a3-1* accumulated higher endogenous ABA levels coupled with a reduced transpiration rate, thereby resulting in a phenotype exhibiting enhanced tolerance to drought stress [77]. The significant increase in *SlNCED1* expression in *ARF4*-as leaves and roots along with the repression of the three ABA catabolism genes in normal and salt stress conditions explain the increase in the concentration of ABA in leaves. This finding suggests the involvement of *ARF4* in the regulation of ABA synthesis and provides clues on the existence of a possible cross talk between auxin and ABA signaling pathways.

In plants, abiotic stresses induce the overproduction of reactive oxygen species (ROS), a highly reactive and toxic molecules that cause damage to proteins, lipids, carbohydrates and DNA and results ultimately in oxidative stress [39]. The ability of plants to mitigate the negative effects of abiotic stresses relays on the efficiency of the antioxidant defense systems to protect plant cells from oxidative damage by scavenging ROS accumulation. The key enzymatic antioxidants are catalase, superoxide dismutase (SOD), monodehydroascorbate reductase (MDHAR), dehydroascorbate reductase (DHAR) [39]. Several studies have linked abiotic stress tolerance to an overproduction of antioxydants in many plant species including tomato [78]. Genetic manipulation of genes encoding for these antioxydants increased plant tolerance to a wide range of abiotic stresses. For instance, the overproduction of a bacterial *cat1* gene improved tolerance to salinity in tomato [79]. Rice plants engineered to express a bacterial *cat1* gene showed increased tolerance to osmotic stress [80]. The overexpression of Rice cytosolic Cu/Zn-SOD in chloroplasts of tobacco plant improved their photosynthetic performance during photooxidative stresses such as high salt or drought stresses [81]. The overexpression of *mdhar* in transgenic tobacco increased the tolerance against salt and osmotic stresses [82]. Here, we found that the expression of *cat1, mdhar* and *SOD* genes was significantly upregulated in *ARF4*-as plants in response to either salt or osmotic stress, which suggest that the down regulation of *ARF4* improved tomato tolerance to salt and osmotic stresses by reducing ROS accumulation mainly through antioxidant enzymes activities.

Genetic diversity in plants is a key source for trait improvement. Creating variations in the gene pool is the main requirement for developing novel plant varieties with desirable traits [83]. Genome-editing technology has revolutionized the world of fundamental and applied research [84]. Highly qualified in terms of versatility, efficiency and specificity, CRISPR technology, the well-known genome-editing technology, had broadened the agricultural research area [85]. In this work, we found that *arf4-cr* mutant showed similar growth and stomatal responses than *ARF4*-as plants. This finding suggests that *arf4-cr* plants may tolerate salt and osmotic stresses.

## 5. Conclusion

This study provides clues on the involvement of *ARF4* in the acquisition of salt and drought tolerance in tomato. We found that the downregulation of *SlARF4* increases tomato tolerance to salinity and drought stress. Indeed, the downregulation of *SlARF4* increases root length and density resulting in a better development of root system. Furthermore, *ARF4*-as plants displayed a leaf curling, a lower stomatal conductance, a higher water use efficiency under normal conditions and maintained high level of chlorophyll content even in stress conditions suggesting that their photosynthetic activity was less affected by the stress application. Total carbohydrates were accumulated in high proportion in photosynthetic tissues of *ARF4*-as plants. At the molecular level, the expression of the sucrose transporter *LeSUT1* was significantly up-regulated in *ARF4*-as which might explain carbohydrates accumulation in root tissues. *cat1, Cu/ZnSOD* and *mdhar* genes were found to be up-regulated in *ARF4*-as plants suggesting that *ARF4*-as mutant is more tolerant to salt and water stress. Furthermore, *SlARF4*-as plants showed a significant increase in ABA content associated with a low stomatal conductance under stressful conditions. This increase in ABA content is due to the activation of ABA biosynthesis genes and the repression of ABA catabolism genes. Besides, *ARF4* mutant generated by CRISPR technology (*arf4-cr*) displayed similar growth and stomatal responses as *ARF4*-as which enable us to confirm the role of *SlARF4*. Taking together, the data presented in this work brings new elements on auxin involvement in stress tolerance in tomato and underline the role of *ARF4* in this process and provides new insights into tomato selection and breeding for environmental stress tolerant.

## Supplementary material

**Figure S1.**
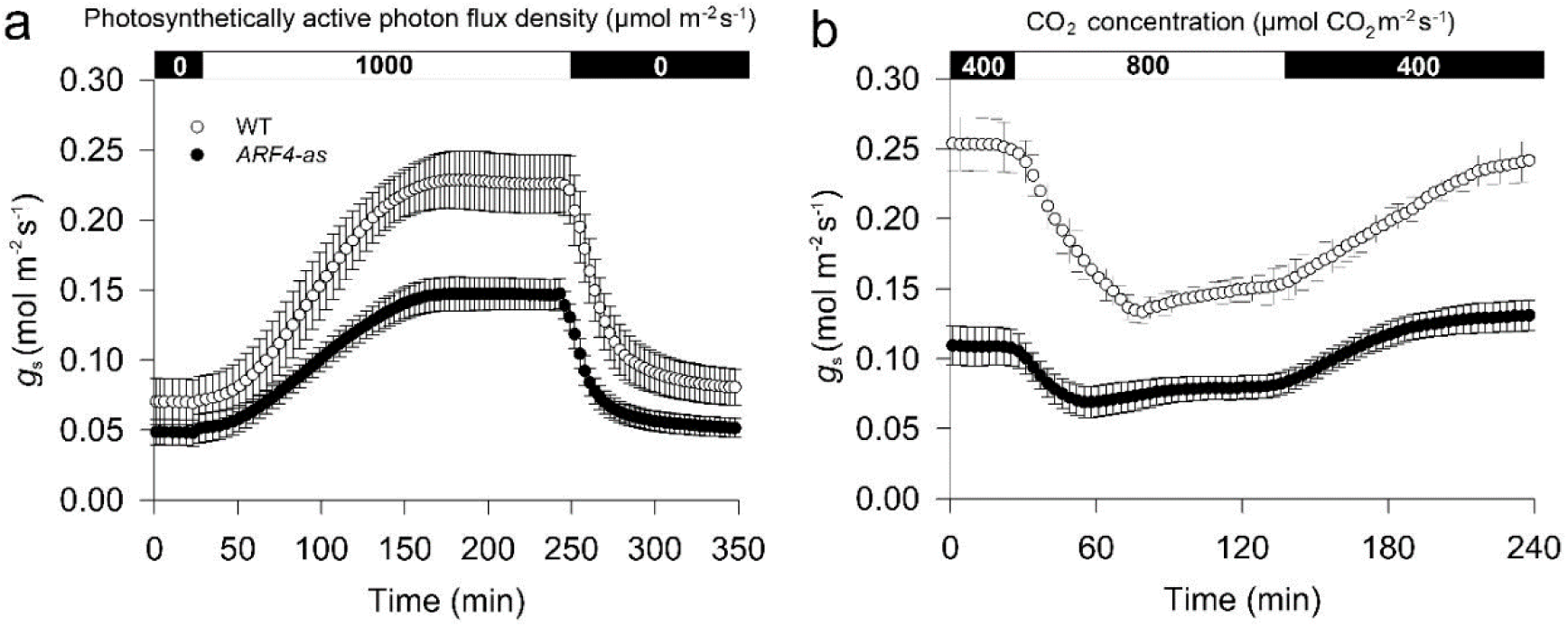
Stomatal responses to irradiance and CO_2_ levels in Micro-Tom (WT) and *ARF4*-as transgenic line. (a) Stomatal conductance (*g*_s_) response in tomato plants cv. Micro-Tom (WT) and isogenic *ARF4* antisense transgenic line (*ARF4-*as) from dark-adapted (0 µmol m^-2^ s^-1^) leaves exposed to light (1000 µmol m^-2^ s^-1^) and on subsequent transfer back to darkness (0 µmol m^-2^ s^-1^) in 350 minutes interval (n=5). (b) Stomatal conductance in response to CO_2_ elevation and subsequent decrease to ambient CO_2_ (400-800-400 µmol CO_2_ m^-2^ s^-1^) in 240 minute interval (n=4). Measurements were performed using a LI-6400; LI-COR gas exchange chamber in plants aged 40 days after germination. Data presented are mean ± SE obtained using the 5^th^ leaf totally expanded.

**Figure S2.**
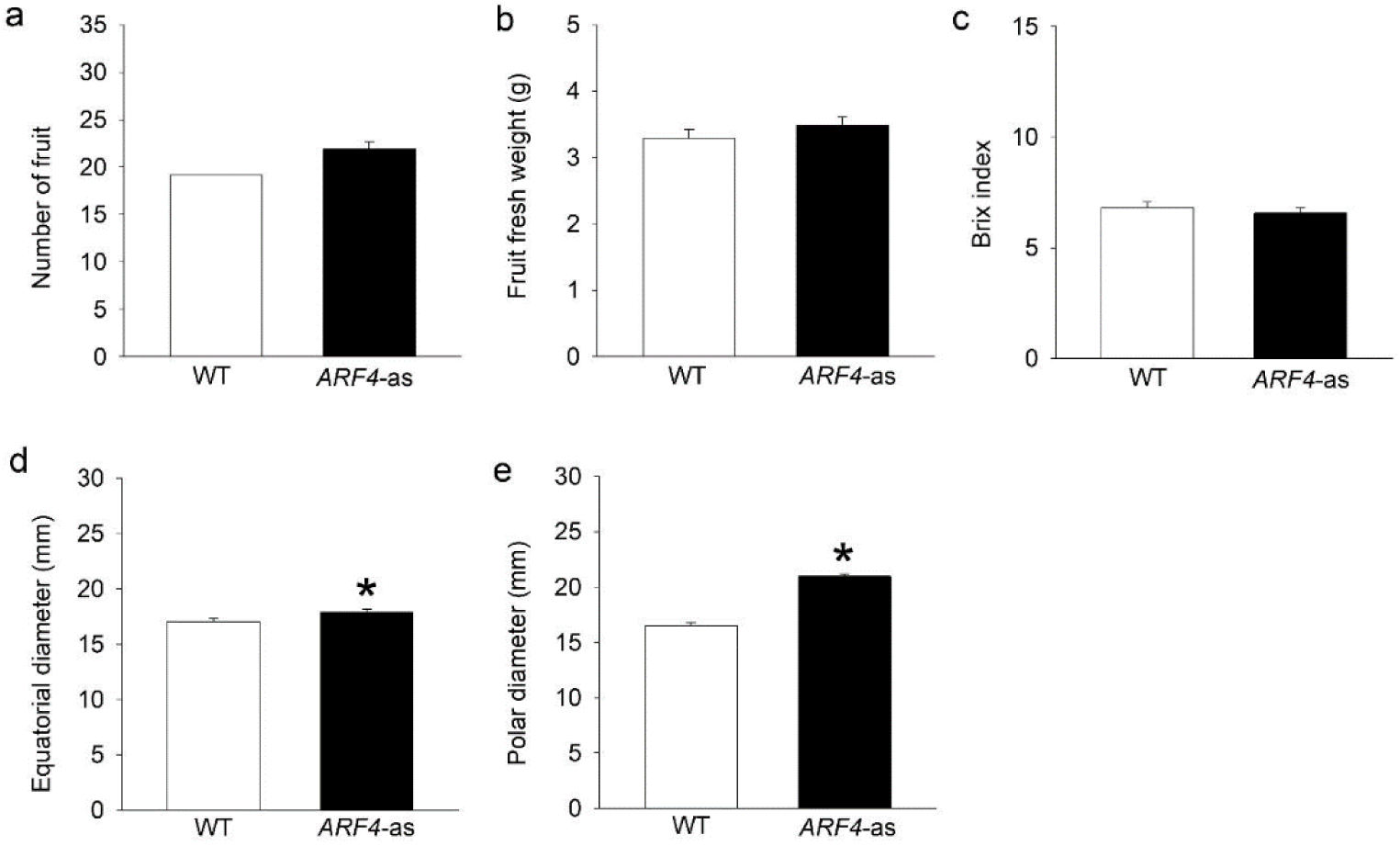
Productive parameters in tomato Micro-Tom (WT) and *ARF4*-as transgenic line. (a) Number of fruit, (b) Fruit fresh weight, (c) brix index, (d) equatorial diameter and (e) polar diameter of the fruit. Values are means ± s.e.m (n=8 plants).Asterisks indicate values that were determined by Student’s t test to be significantly different (P < 0.05) from Micro-tom (WT).

**Figure S3.**
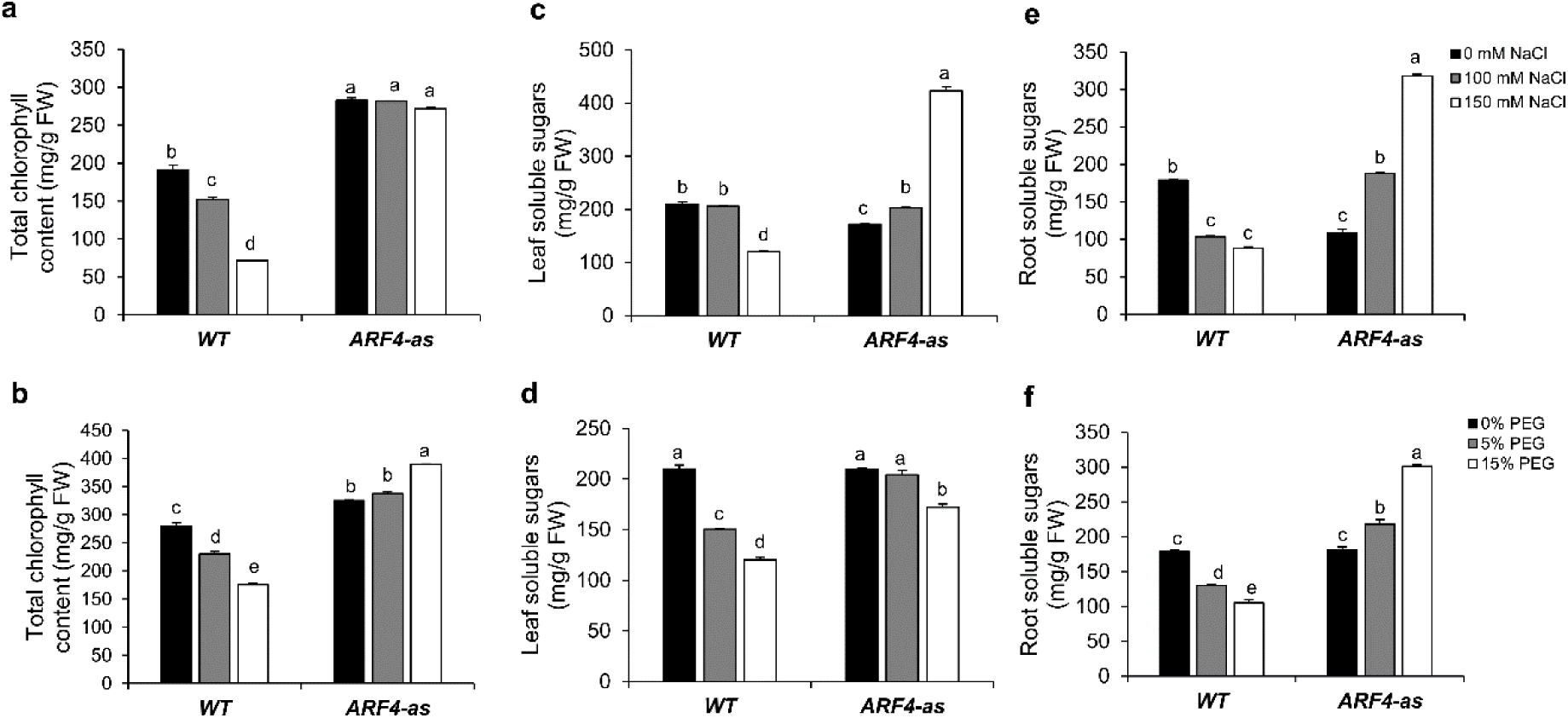
Photosynthesis and sugar accumulation in WT and *ARF4*-as plants exposed to different concentrations of NaCl or PEG. (a) and (b) total chlorophyll content in salt and osmotic stress conditions respectively, (c) and (d) leaf soluble sugars in salt and osmotic stress conditions respectively, (e) and (f) root soluble sugars content in salt and osmotic stress conditions respectively. Salt and osmotic stresses were performed on three weeks tomato plants for two weeks by adding 100mM of NaCl or 150 mM of NaCl for salt stress or 5% or 15% of PEG 20 000 for osmotic stress. Values are mean ± SD of three biological replicates. Bars with different letters indicate the statistical significance (p<0,05) according to Student Newman-Keuls test.

**Figure S4.**
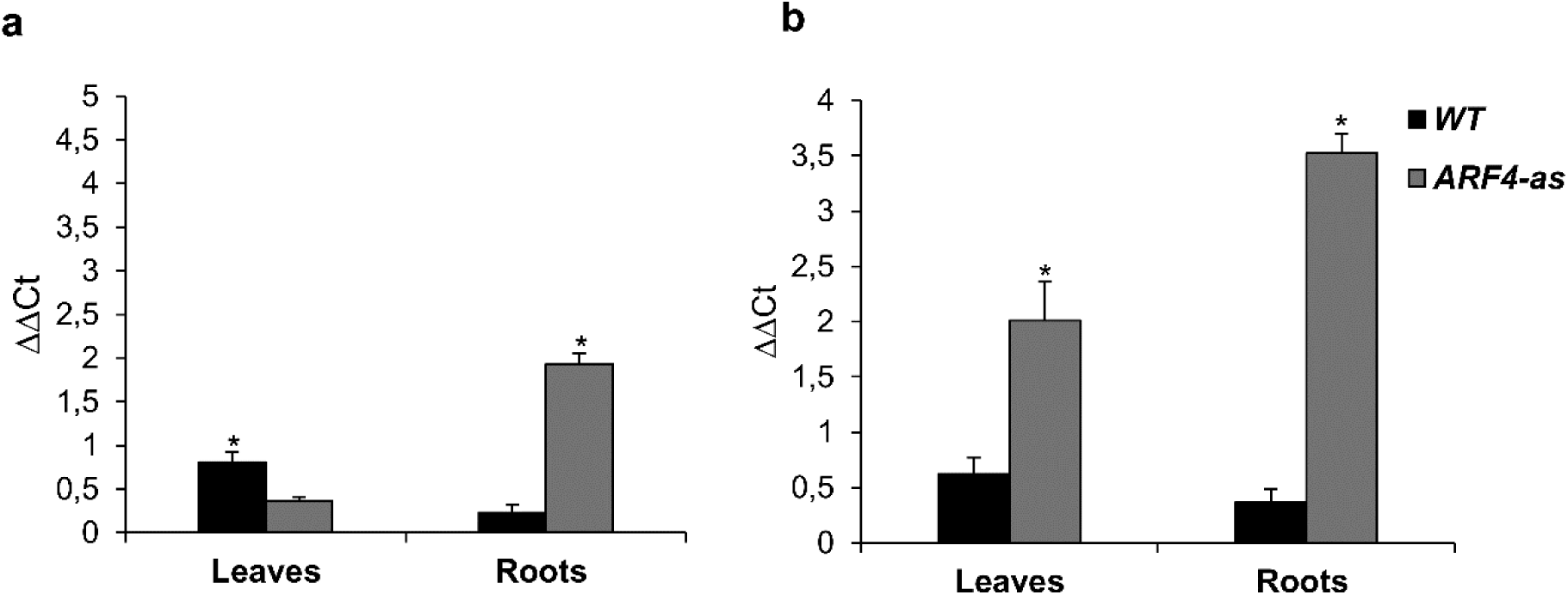
Expression of sucrose transporter *LeSUT1* in WT and *ARF4*-as plants exposed to salt or osmotic stresses. (a) gene expression in leaves and roots exposed to 150mM of NaCl, (b) gene expression in leaves and roots leaves exposed to 15% PEG. ΔΔCt refers to fold differences in gene expression relative to untreated plants. Values are mean ± SD of three biological replicates. Stars (*) indicate the statistical significance (p<0,05) according to Student’s t-test.

**Figure S5.**
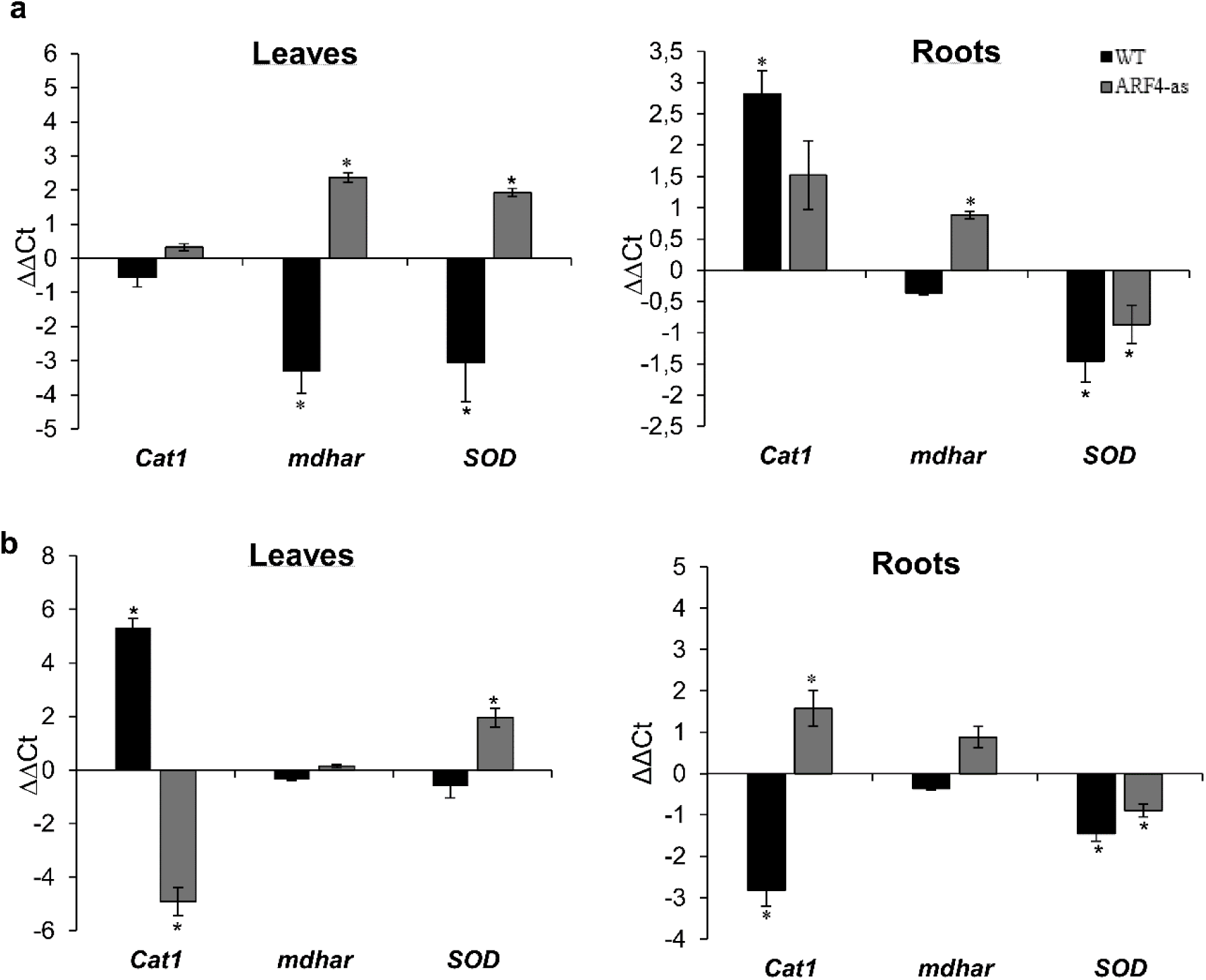
Expression of *Cat1, mdhar* and *SOD* in WT and *ARF4*-as plants exposed to salt or osmotic stresses. (a) gene expression in leaves and roots exposed to 150mM of NaCl, (b) gene expression in leaves and roots leaves exposed to 15% PEG. ΔΔCt refers to fold differences in gene expression relative to untreated plants. Values are mean ± SD of three biological replicates. Stars (*) indicate the statistical significance (p<0,05) using Student’s t-test.

**Figure S6.**
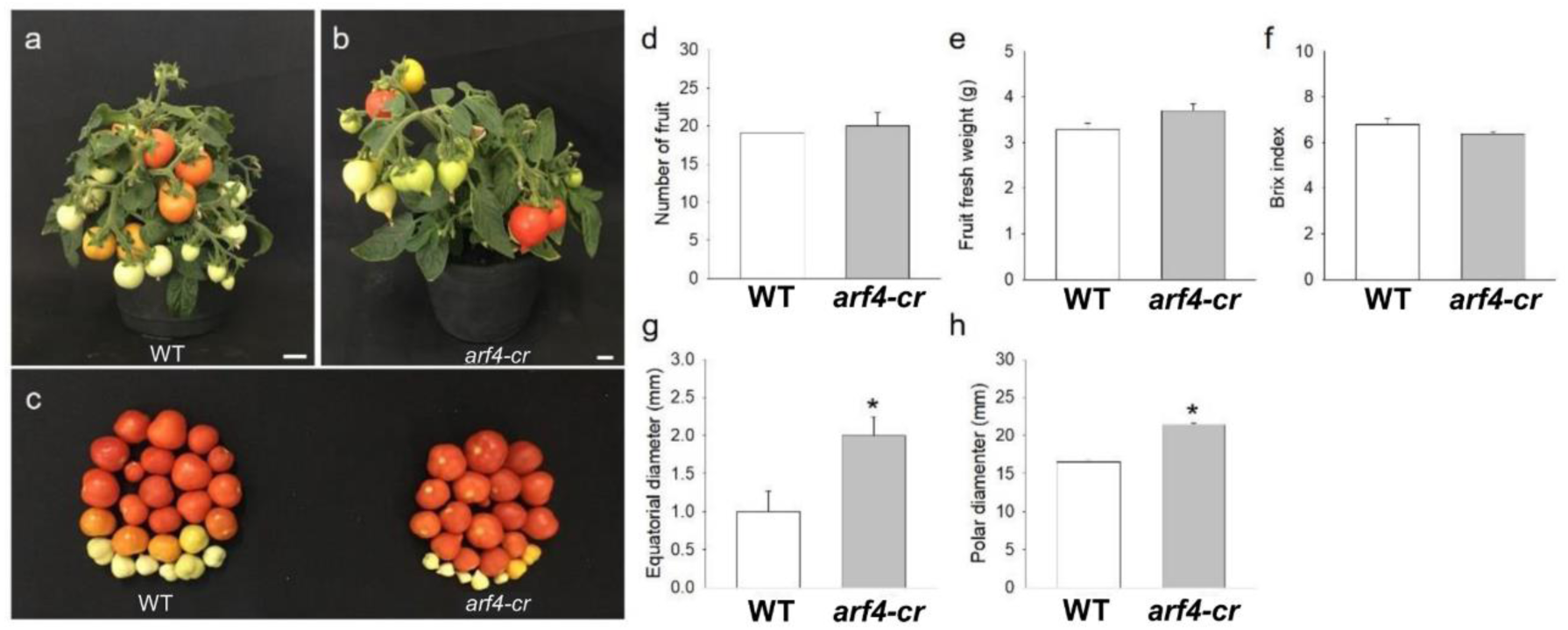
Productive parameters in tomato Micro-Tom (WT) and *arf4-cr* transgenic line. (a) and (b) representative tomato plants in reproductive stage, (c) fruit yield, (d) Number of fruit, (e) Fruit fresh weight, (f) percentage of soluble solids, (g) Equatorial diameter of the fruit and (h) Polar diameter of the fruit. Values are means ± s.e.m (n=8). The brix index was measured in 10 fruits per repetition. Asterisks indicate values that were determined by Student’s t test to be significantly different (P < 0.05) from wild-type (WT).

**Table S1.**
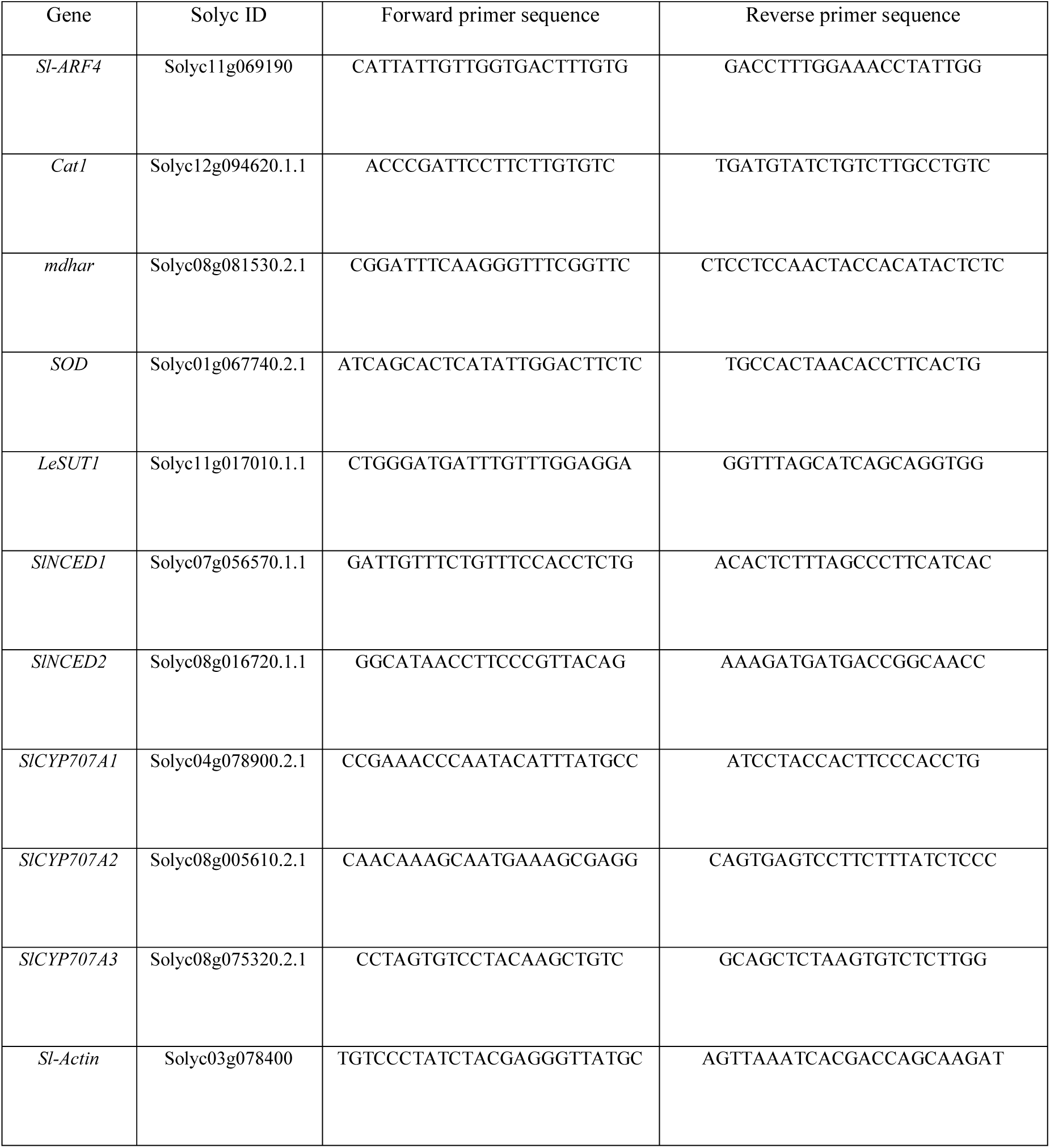
Gene ID and quantitative RT-PCR primers.

**Table S2.** Characterization of photosynthetic parameters in tomato cv. Micro-Tom (WT) and isogenic *ARF4* antisense transgenic line (*ARF4-*as) Vcmax: maximum Rubisco carboxylation rate; Jmax: maximum electron transport rate; TPU: triose phosphate utilization. Mean values (n=5) ± s.e.m. Significant differences by t-test at 0.01.

|  | WT | *ARF4*-as |
| --- | --- | --- |
| $V_{\text{cmax}}$ ($\mu\text{mol m}^{-2} \text{s}^{-1}$) | 80.37 ± 2.34 | 70.43* ± 3.01 |
| $J_{\text{max}}$ ($\mu\text{mol m}^{-2} \text{s}^{-1}$) | 156.40 ± 7.74 | 140.70* ± 6.30 |
| TPU ($\mu\text{mol m}^{-2} \text{s}^{-1}$) | 11.84 ± 0.57 | 10.82 ± 0.53 |

## Author contributions

All authors have read and agree to the published version of the manuscript. Conceptualization, N.B., M.B, A.Z., A.S. and M.Z.; methodology, A.Z., A.S. and M.Z.; formal analysis, S.B., K.G., G.H., M.M.B., B.L.; writing original draft preparation, S.B., A.Z., K.G, A.S. and M.Z.; writing—review and editing, N.B., M.B., F.M., A.Z., A.S. and M.Z.; supervision, A.Z., A.S. and M.Z.; funding acquisition, M.Z.

## Acknowledgments

The authors are grateful to L. Lemonnier, D. Saint-Martin and O. Berseille for genetic transformation and culture of tomato plants and I. Mila for the cloning.

## References

1. Vanneste, S.; Friml, J. Auxin: a trigger for change in plant development. Cell 2009, 136, 1005–1016.

2. Szemenyei, H.; Hannon, M.; Long, J.A. TOPLESS mediates auxin-dependent transcriptional repression during Arabidopsis embryogenesis. Science 2008, 319, 1384–1386.

3. Guilfoyle, T.J.; Hagen, G. Auxin response factors. Curr. Opin. Plant Biol. 2007, 10, 453–460.

4. Zouine, M.; Fu, Y.; Chateigner-Boutin, A.-L.; Mila, I.; Frasse, P.; Wang, H.; Audran, C.; Roustan, J.-P.; Bouzayen, M. Characterization of the tomato ARF gene family uncovers a multi-levels post-transcriptional regulation including alternative splicing. PloS One 2014, 9, e84203.

5. Ulmasov, T.; Murfett, J.; Hagen, G.; Guilfoyle, T.J. Aux/IAA proteins repress expression of reporter genes containing natural and highly active synthetic auxin response elements. Plant Cell 1997, 9, 1963–1971.

6. Jain, M.; Khurana, J.P. Transcript profiling reveals diverse roles of auxin-responsive genes during reproductive development and abiotic stress in rice. FEBS J. 2009, 276, 3148–3162.

7. Xing, H.; Pudake, R.N.; Guo, G.; Xing, G.; Hu, Z.; Zhang, Y.; Sun, Q.; Ni, Z. Genome-wide identification and expression profiling of auxin response factor (ARF) gene family in maize. BMC Genomics 2011, 12, 178.

8. Hu, W.; Zuo, J.; Hou, X.; Yan, Y.; Wei, Y.; Liu, J.; Li, M.; Xu, B.; Jin, Z. The auxin response factor gene family in banana: genome-wide identification and expression analyses during development, ripening, and abiotic stress. Crop Sci. Hortic. 2015, 742.

9. Tang, Y.; Bao, X.; Liu, K.; Wang, J.; Zhang, J.; Feng, Y.; Wang, Y.; Lin, L.; Feng, J.; Li, C. Genome-wide identification and expression profiling of the auxin response factor (ARF) gene family in physic nut. PloS One 2018, 13, e0201024.

10. Zhang, X.; Yan, F.; Tang, Y.; Yuan, Y.; Deng, W.; Li, Z. Auxin response gene SlARF3 plays multiple roles in tomato development and is involved in the formation of epidermal cells and trichomes. Plant Cell Physiol. 2015, 56, 2110–2124.

11. Breitel, D.A.; Chappell-Maor, L.; Meir, S.; Panizel, I.; Puig, C.P.; Hao, Y.; Yifhar, T.; Yasuor, H.; Zouine, M.; Bouzayen, M.; et al. AUXIN RESPONSE FACTOR 2 Intersects Hormonal Signals in the Regulation of Tomato Fruit Ripening. PLOS Genet. 2016, 12, e1005903.

12. Liu, N.; Wu, S.; Houten, J. Van; Wang, Y.; Ding, B.; Fei, Z.; Clarke, T.H.; Reed, J.W.; Van Der Knaap, E. Down-regulation of AUXIN RESPONSE FACTORS 6 and 8 by microRNA 167 leads to floral development defects and female sterility in tomato. J. Exp. Bot. 2014, 65, 2507–2520.

13. Jones, B.; Frasse, P.; Olmos, E.; Zegzouti, H.; Li, Z.G.; Latché, A.; Pech, J.C.; Bouzayen, M. Down-regulation of DR12, an auxin-response-factor homolog, in the tomato results in a pleiotropic phenotype including dark green and blotchy ripening fruit. Plant J. 2002, 32, 603–613.

14. Sagar, M.; Chervin, C.; Mila, I.; Hao, Y.; Roustan, J.-P.; Benichou, M.; Gibon, Y.; Biais, B.; Maury, P.; Latché, A.; et al. SlARF4, an auxin response factor involved in the control of sugar metabolism during tomato fruit development. Plant Physiol. 2013, 161, 1362–74.

15. Mei, L.; Yuan, Y.; Wu, M.; Gong, Z.; Zhang, Q.; Yang, F.; Zhang, Q.; Luo, Y.; Xu, X.; Zhang, W. SlARF10, an auxin response factor, is required for chlorophyll and sugar accumulation during tomato fruit development. bioRxiv 2018, 253237.

16. Bouzroud, S.; Gouiaa, S.; Hu, N.; Bernadac, A.; Mila, I.; Bendaou, N.; Smouni, A.; Bouzayen, M.; Zouine, M. Auxin Response Factors (ARFs) are potential mediators of auxin action in tomato response to biotic and abiotic stress (Solanum lycopersicum). PLOS ONE 2018, 13, e0193517.

17. Wang, H.; Jones, B.; Li, Z.; Frasse, P.; Delalande, C.; Regad, F.; Chaabouni, S.; Latché, A.; Pech, J.-C.; Bouzayen, M. The tomato Aux/IAA transcription factor IAA9 is involved in fruit development and leaf morphogenesis. Plant Cell 2005, 17, 2676–2692.

18. Brooks, C.; Nekrasov, V.; Lippman, Z.B.; Van Eck, J. Efficient gene editing in tomato in the first generation using the clustered regularly interspaced short palindromic repeats/CRISPR-associated9 system. Plant Physiol. 2014, 166, 1292–1297.

19. Lei, Y.; Lu, L.; Liu, H.-Y.; Li, S.; Xing, F.; Chen, L.-L. CRISPR-P: a web tool for synthetic single-guide RNA design of CRISPR-system in plants. Mol. Plant 2014, 7, 1494–1496.

20. Hunt, R. Plant Growth Analysis: Second derivatives and compounded second derivatives of splined plant growth curves. Ann. Bot. 1982, 50, 317–328.

21. Farquhar, G.D.; von Caemmerer, S.; Berry, J.A. A biochemical model of photosynthetic CO2 assimilation in leaves of C3 species. Planta 1980, 149, 78–90.

22. Rodeghiero, M.; Niinemets, Ü.; Cescatti, A. Major diffusion leaks of clamp-on leaf cuvettes still unaccounted: How erroneous are the estimates of Farquhar et al. model parameters? Plant Cell Environ. 2007, 30, 1006–1022.

23. Broughton, W.J.; Dilworth, M.J. Control of leghaemoglobin synthesis in snake beans. Biochem. J. 1971, 125, 1075–1080.

24. Bassa, C.; Mila, I.; Bouzayen, M.; Audran-Delalande, C. Phenotypes associated with down-regulation of Sl-IAA27 support functional diversity among Aux/IAA family members in the tomato. Plant Cell Physiol. 2012, pcs101.

25. Dubois, M.; Gilles, K.A.; Hamilton, J.K.; Rebers, P. t; Smith, F. Colorimetric method for determination of sugars and related substances. Anal. Chem. 1956, 28, 350–356.

26. Forcat, S.; Bennett, M.H.; Mansfield, J.W.; Grant, M.R. A rapid and robust method for simultaneously measuring changes in the phytohormones ABA, JA and SA in plants following biotic and abiotic stress. Plant Methods 2008, 4, 16.

27. Jaulneau, V.; Lafitte, C.; Jacquet, C.; Fournier, S.; Salamagne, S.; Briand, X.; Esquerré-Tugayé, M.-T.; Dumas, B. Ulvan, a Sulfated Polysaccharide from Green Algae, Activates Plant Immunity through the Jasmonic Acid Signaling Pathway. J. Biomed. Biotechnol. 2010, 2010.

28. Smart, R.E.; Bingham, G.E. Rapid Estimates of Relative Water Content. Plant Physiol. 1974, 53, 258–260.

29. Bernacchi, C.J.; Rosenthal, D.M.; Pimentel, C.; Long, S.P.; Farquhar, G.D. Modeling the temperature dependence of C 3 photosynthesis. In Photosynthesis in silico; Springer, 2009; pp. 231–246.

30. Jiang, Q.; Roche, D.; Monaco, T.; Hole, D. Stomatal conductance is a key parameter to assess limitations to photosynthesis and growth potential in barley genotypes. Plant Biol. 2006, 8, 515–521.

31. Ashraf, M.; Harris, P. Photosynthesis under stressful environments: an overview. Photosynthetica 2013, 51, 163–190.

32. De Jong, M.; Mariani, C.; Vriezen, W.H. The role of auxin and gibberellin in tomato fruit set. J. Exp. Bot. 2009, 60, 1523–1532.

33. Munns, R.; James, R.A.; Läuchli, A. Approaches to increasing the salt tolerance of wheat and other cereals. J. Exp. Bot. 2006, 57, 1025–1043.

34. Jia, W.; Zhang, L.; Wu, D.; Liu, S.; Gong, X.; Cui, Z.; Cui, N.; Cao, H.; Rao, L.; Wang, C. Sucrose transporter AtSUC9 mediated by a low sucrose level is involved in Arabidopsis abiotic stress resistance by regulating sucrose distribution and ABA accumulation. Plant Cell Physiol. 2015, 56, 1574–1587.

35. Damour, G.; Simonneau, T.; Cochard, H.; Urban, L. An overview of models of stomatal conductance at the leaf level. Plant Cell Environ. 2010, 33, 1419–38.

36. Schultz, H.R. Differences in hydraulic architecture account for near-isohydric and anisohydric behaviour of two field-grown Vitis vinifera L. cultivars during drought. Plant Cell Environ. 2003, 26, 1393–1405.

37. Leymarie, J.; Lascève, G.; Vavasseur, A. Interaction of stomatal responses to ABA and CO2 in Arabidopsis thaliana. Funct. Plant Biol. 1998, 25, 785–791.

38. Yang, R.; Yang, T.; Zhang, H.; Qi, Y.; Xing, Y.; Zhang, N.; Li, R.; Weeda, S.; Ren, S.; Ouyang, B.; et al. Hormone profiling and transcription analysis reveal a major role of ABA in tomato salt tolerance. Plant Physiol. Biochem. 2014, 77, 23–34.

39. Das, K.; Roychoudhury, A. Reactive oxygen species (ROS) and response of antioxidants as ROS-scavengers during environmental stress in plants. Environ. Toxicol. 2014, 2, 53.

40. Choudhury, S.; Panda, P.; Sahoo, L.; Panda, S.K. Reactive oxygen species signaling in plants under abiotic stress. Plant Signal. Behav. 2013, 8, e23681.

41. Marin, E.; Jouannet, V.; Herz, A.; Lokerse, A.S.; Weijers, D.; Vaucheret, H.; Nussaume, L.; Crespi, M.D.; Maizel, A. miR390, Arabidopsis TAS3 tasiRNAs, and their AUXIN RESPONSE FACTOR targets define an autoregulatory network quantitatively regulating lateral root growth. Plant Cell 2010, 22, 1104–1117.

42. Pereira-Netto, A.B. de; Gabriele, A.C.; Pinto, H.S. Aspects of leaf anatomy of kudzu (Pueraria lobata, Leguminosae-Faboideae) related to water and energy balance. Pesqui. Agropecu. Bras. 1999, 34, 1361–1365.

43. Jones, B.; Frasse, P.; Olmos, E.; Zegzouti, H.; Li, Z.G.; Latché, A.; Pech, J.C.; Bouzayen, M. Down-regulation of DR12, an auxin-response-factor homolog, in the tomato results in a pleiotropic phenotype including dark green and blotchy ripening fruit. Plant J. 2002, 32, 603–613.

44. Kadioglu, A.; Terzi, R.; Saruhan, N.; Saglam, A. Current advances in the investigation of leaf rolling caused by biotic and abiotic stress factors. Plant Sci. 2012, 182, 42–8.

45. Fen, L.L.; Ismail, M.R.; Zulkarami, B.; Rahman, M.S.A.; Islam, M.R. Physiological and molecular characterization of drought responses and screening of drought tolerant rice varieties. Biosci. J. 2015, 31.

46. Zhang, Y.; Wang, Z.; Wu, Y.; Zhang, X. Stomatal characteristics of different green organs in wheat under different irrigation regimes. Zuo Wu Xue Bao 2006, 32, 70–75.

47. Brugnoli, E.; Lauteri, M. Effects of Salinity on Stomatal Conductance, Photosynthetic Capacity, and Carbon Isotope Discrimination of Salt-Tolerant (Gossypium hirsutum L.) and Salt-Sensitive (Phaseolus vulgaris L.) C3 Non-Halophytes. Plant Physiol. 1991, 95, 628–635.

48. Mafakheri, A.; Siosemardeh, A.; Bahramnejad, B.; Struik, P.; Sohrabi, Y. Effect of drought stress on yield, proline and chlorophyll contents in three chickpea cultivars. Aust. J. Crop Sci. 2010, 4, 580.

49. Rahman, A. Auxin: a regulator of cold stress response. Physiol. Plant. 2013, 147, 28–35.

50. Navarro, L.; Dunoyer, P.; Jay, F.; Arnold, B.; Dharmasiri, N.; Estelle, M.; Voinnet, O.; Jones, J.D. A plant miRNA contributes to antibacterial resistance by repressing auxin signaling. Science 2006, 312, 436–439.

51. Guilfoyle, T.J.; Hagen, G. Auxin response factors. Curr. Opin. Plant Biol. 2007, 10, 453–460.

52. Wang, S.; Bai, Y.; Shen, C.; Wu, Y.; Zhang, S.; Jiang, D.; Guilfoyle, T.J.; Chen, M.; Qi, Y. Auxin-related gene families in abiotic stress response in Sorghum bicolor. Funct. Integr. Genomics 2010, 10, 533–546.

53. Van Ha, C.; Le, D.T.; Nishiyama, R.; Watanabe, Y.; Sulieman, S.; Tran, U.T.; Mochida, K.; Van Dong, N.; Yamaguchi-Shinozaki, K.; Shinozaki, K. The auxin response factor transcription factor family in soybean: genome-wide identification and expression analyses during development and water stress. DNA Res. 2013, 20, 511–524.

54. Guóth, A.; Tari, I.; Gallé, Á.; Csiszár, J.; Pécsváradi, A.; Cseuz, L.; Erdei, L. Comparison of the drought stress responses of tolerant and sensitive wheat cultivars during grain filling: changes in flag leaf photosynthetic activity, ABA levels, and grain yield. J. Plant Growth Regul. 2009, 28, 167–176.

55. Farooq, M.; Wahid, A.; Kobayashi, N.; Fujita, D.; Basra, S. Plant drought stress: effects, mechanisms and management. In Sustainable agriculture; Springer, 2009; pp. 153–188.

56. Saddam, S.; Bibi, A.; Sadaqat, H.; Usman, B. Comparison of 10 sorghum (Sorghum bicolor L.) genotypes under various water stress regimes. J Anim Plant Sci 2014, 24, 1811–1820.

57. Fayez, K.A.; Bazaid, S.A. Improving drought and salinity tolerance in barley by application of salicylic acid and potassium nitrate. J. Saudi Soc. Agric. Sci. 2014, 13, 45–55.

58. Akinci, Ş.; Lösel, D.M. Plant water-stress response mechanisms. In Water stress; InTech, 2012.

59. Kurth, E.; Cramer, G.R.; Läuchli, A.; Epstein, E. Effects of NaCl and CaCl2 on Cell Enlargement and Cell Production in Cotton Roots. Plant Physiol. 1986, 82, 1102–1106.

60. Burssens, S.; Himanen, K.; van de Cotte, B.; Beeckman, T.; Van Montagu, M.; Inze, D.; Verbruggen, N. Expression of cell cycle regulatory genes and morphological alterations in response to salt stress in Arabidopsis thaliana. Planta 2000, 211, 632–640.

61. Comas, L.H.; Becker, S.R.; Cruz, V.M. V.; Byrne, P.F.; Dierig, D.A. Root traits contributing to plant productivity under drought. Front. Plant Sci. 2013, 4.

62. Munns, R.; James, R.A.; Läuchli, A. Approaches to increasing the salt tolerance of wheat and other cereals. J. Exp. Bot. 2006, 57, 1025–1043.

63. Munns, R.; James, R.A.; Läuchli, A. Approaches to increasing the salt tolerance of wheat and other cereals. J. Exp. Bot. 2006, 57, 1025–1043.

64. Dogan, M.; Tipirdamaz, R.; Demir, Y. Salt resistance of tomato species grown in sand culture. Plant Soil Env. 2010, 56, 499–507.

65. Zarafshar, M.; Akbarinia, M.; Askari, H.; Mohsen Hosseini, S.; Rahaie, M.; Struve, D.; Gabriel Striker, G. Morphological, physiological and biochemical responses to soil water deficit in seedlings of three populations of wild pear tree (Pyrus boisseriana). Base 2014.

66. Hernandez, J.; Jimenez, A.; Mullineaux, P.; Sevilia, F. Tolerance of pea (Pisum sativum L.) to long-term salt stress is associated with induction of antioxidant defences. Plant Cell Environ. 2000, 23, 853–862.

67. Hasegawa, P.M.; Bressan, R.A.; Zhu, J.-K.; Bohnert, H.J. Plant cellular and molecular responses to high salinity. Annu. Rev. Plant Biol. 2000, 51, 463–499.

68. Atkinson, N.J.; Urwin, P.E. The interaction of plant biotic and abiotic stresses: from genes to the field. J. Exp. Bot. 2012, ers100.

69. Patade, V.Y.; Bhargava, S.; Suprasanna, P. Transcript expression profiling of stress responsive genes in response to short-term salt or PEG stress in sugarcane leaves. Mol. Biol. Rep. 2012, 39, 3311–3318.

70. Xie, C.; Zhang, R.; Qu, Y.; Miao, Z.; Zhang, Y.; Shen, X.; Wang, T.; Dong, J. Overexpression of MtCAS31 enhances drought tolerance in transgenic Arabidopsis by reducing stomatal density. New Phytol. 2012, 195, 124–135.

71. Li, J.; Li, Y.; Yin, Z.; Jiang, J.; Zhang, M.; Guo, X.; Ye, Z.; Zhao, Y.; Xiong, H.; Zhang, Z. Os ASR 5 enhances drought tolerance through a stomatal closure pathway associated with ABA and H2O2 signalling in rice. Plant Biotechnol. J. 2017, 15, 183–196.

72. Lee, S.C.; Lim, C.W.; Lan, W.; He, K.; Luan, S. ABA signaling in guard cells entails a dynamic protein–protein interaction relay from the PYL-RCAR family receptors to ion channels. Mol. Plant 2013, 6, 528–538.

73. Schroeder, J.I.; Kwak, J.M.; Allen, G.J. Guard cell abscisic acid signalling and engineering drought hardiness in plants. Nature 2001, 410, 327.

74. Schroeder, J.I.; Allen, G.J.; Hugouvieux, V.; Kwak, J.M.; Waner, D. Guard cell signal transduction. Annu. Rev. Plant Biol. 2001, 52, 627–658.

75. Burbidge, A.; Grieve, T.M.; Jackson, A.; Thompson, A.; McCarty, D.R.; Taylor, I.B. Characterization of the ABA-deficient tomato mutant notabilis and its relationship with maize Vp14. Plant J. 1999, 17, 427–431.

76. Wan, X.-R.; Li, L. Regulation of ABA level and water-stress tolerance of Arabidopsis by ectopic expression of a peanut 9-cis-epoxycarotenoid dioxygenase gene. Biochem. Biophys. Res. Commun. 2006, 347, 1030–1038.

77. Umezawa, T.; Okamoto, M.; Kushiro, T.; Nambara, E.; Oono, Y.; Seki, M.; Kobayashi, M.; Koshiba, T.; Kamiya, Y.; Shinozaki, K. CYP707A3, a major ABA 8′-hydroxylase involved in dehydration and rehydration response in Arabidopsis thaliana. Plant J. 2006, 46, 171–182.

78. Gondim, F.A.; Gomes-Filho, E.; Costa, J.H.; Mendes Alencar, N.L.; Prisco, J.T. Catalase plays a key role in salt stress acclimation induced by hydrogen peroxide pretreatment in maize. Plant Physiol. Biochem. PPB Société Fr. Physiol. Végétale 2012, 56, 62–71.

79. Mohamed, E.-A.; Iwaki, T.; Munir, I.; Tamoi, M.; Shigeoka, S.; Wadano, A. Overexpression of bacterial catalase in tomato leaf chloroplasts enhances photo-oxidative stress tolerance. Plant Cell Environ. 2003, 26, 2037–2046.

80. Moriwaki, T.; Yamamoto, Y.; Aida, T.; Funahashi, T.; Shishido, T.; Asada, M.; Prodhan, S.H.; Komamine, A.; Motohashi, T. Overexpression of the Escherichia coli catalase gene, katE, enhances tolerance to salinity stress in the transgenic indica rice cultivar, BR5. Plant Biotechnol. Rep. 2008, 2, 41–46.

81. Badawi, G.H.; Yamauchi, Y.; Shimada, E.; Sasaki, R.; Kawano, N.; Tanaka, K.; Tanaka, K. Enhanced tolerance to salt stress and water deficit by overexpressing superoxide dismutase in tobacco (Nicotiana tabacum) chloroplasts. Plant Sci. 2004, 166, 919–928.

82. Eltayeb, A.E.; Kawano, N.; Badawi, G.H.; Kaminaka, H.; Sanekata, T.; Shibahara, T.; Inanaga, S.; Tanaka, K. Overexpression of monodehydroascorbate reductase in transgenic tobacco confers enhanced tolerance to ozone, salt and polyethylene glycol stresses. Planta 2006, 225, 1255–1264.

83. Arora, L.; Narula, A. Gene Editing and Crop Improvement Using CRISPR-Cas9 System. Front. Plant Sci. 2017, 8.

84. Xu, J.; Hua, K.; Lang, Z. Genome editing for horticultural crop improvement. Hortic. Res. 2019, 6, 1–16.

85. Bortesi, L.; Fischer, R. The CRISPR/Cas9 system for plant genome editing and beyond. Biotechnol. Adv. 2015, 33, 41–52.

